# Stretch Regulates Alveologenesis and Homeostasis Via Mesenchymal G_αq/11_-Mediated TGFβ2 Activation

**DOI:** 10.1101/2020.09.06.284778

**Authors:** Amanda T Goodwin, Alison E John, Chitra Joseph, Anthony Habgood, Amanda L Tatler, Katalin Susztak, Matthew Palmer, Stefan Offermanns, Neil C Henderson, R Gisli Jenkins

## Abstract

Alveolar development and repair require tight spatiotemporal regulation of numerous signalling pathways that are influenced by chemical and mechanical stimuli. Mesenchymal cells play key roles in numerous developmental processes. Transforming growth factor-β (TGFβ) is essential for alveologenesis and lung repair, and the G protein α subunits G_αq_ and G_α11_ (G_αq/11_) transmit mechanical and chemical signals to activate TGFβ in epithelial cells. To understand the role of mesenchymal G_αq/11_ in lung development, we generated constitutive (*Pdgfrb-Cre^+/−^;Gnaq^fl/fl^;Gna11^−/−^*) and inducible (*Pdgfrb-Cre/ERT2^+/−^;Gnaq^fl/fl^;Gna11^−/−^*) mesenchymal G_αq/11_ deleted mice. Mice with constitutive G_αq/11_ gene deletion exhibited abnormal alveolar development, with suppressed myofibroblast differentiation, altered mesenchymal cell synthetic function, and reduced lung TGFβ2 deposition, as well as kidney abnormalities. Tamoxifen-induced mesenchymal G_αq/11_ gene deletion in adult mice resulted in emphysema associated with reduced TGFβ2 and elastin deposition. Cyclical mechanical stretch-induced TGFβ activation required G_αq/11_ signalling and serine protease activity, but was independent of integrins, suggesting an isoform-specific role for TGFβ2. These data highlight a previously undescribed mechanism of cyclical stretch-induced G_αq/11_-dependent TGFβ2 signalling in mesenchymal cells, which is imperative for normal alveologenesis and maintenance of lung homeostasis.

**Summary statement:** Mesenchymal cell G_αq/11_ signalling regulates myofibroblast function and stretch-mediated TGFβ2 signalling, which are important for alveologenesis and organ homeostasis. These mechanisms are relevant to both developmental and adult lung disease.

## Introduction

Normal alveologenesis requires tight spatiotemporal control of numerous molecular signalling pathways, and coordinated crosstalk between multiple cell types. Any perturbation to these complex processes can disrupt alveolar formation, resulting in structural and functional abnormalities to the gas exchange regions of the lungs. Such abnormalities contribute to perinatal death and lifelong lung function disturbances in survivors (Lovering et al. 2014). The alveolar stage is the final phase of lung development, during which primitive pulmonary sacculi are divided by newly formed secondary septae to form mature alveoli. Alveolarisation occurs between 36 weeks gestation to around 6 years of age in humans (Donahoe, Longoni, and High 2016), and postnatal days 3-30 (P3-P30) in mice (Beauchemin et al. 2016; Pozarska et al. 2017; C. Li et al. 2015), therefore postnatal exposures and stimuli are key influences in alveolar development. Many pathways that drive normal lung development are also instrumental in adult lung repair (Chanda et al. 2019), therefore understanding normal lung development could have implications for numerous pulmonary diseases.

Pericytes are perivascular cells that are widely considered to be mesenchymal precursor cells in the lung, and are integral to multiple developmental processes (Kato et al. 2018; Barron, Gharib, and Duffield 2016; Ricard et al. 2014). Pericytes express platelet-derived growth factor-β (PDGFRβ), PDGFRα, and NG2, among other markers, but the most specific marker for pericytes is PDGFRβ (Riccetti et al. 2020). Pericytes migrate and differentiate into parenchymal myofibroblasts in the lung, and myofibroblast-driven deposition of extracellular matrix (ECM) proteins, such as collagen and elastin, provide the essential scaffolds for secondary septation during lung development and lung repair (Mecham 2018; Mizikova and Morty 2015). Therefore pericytes, and the mesenchymal cells that derive from them, are instrumental in alveologenesis and lung homeostasis.

The pleiotropic cytokine transforming growth factor-β (TGFβ) regulates numerous developmental and repair processes, including the proliferation, migration, and differentiation of pericytes (Bartram and Speer 2004), and the generation of ECM. TGFβ signalling is tightly regulated in vivo by the production of TGFβ in latent form, and the three mammalian TGFβ isoforms must be activated to exert their biological effects. While TGFβ signalling is essential for multiple processes in alveolar development and repair (Bartram and Speer 2004), the mechanisms that control TGFβ activation in alveologenesis are unclear.

Latent TGFβ is activated when a conformational change to the large latent complex alters the relationship between TGFβ and the latency associated peptide, allowing TGFβ to interact with its receptor. The G-proteins G_αq_ and G_α11_ (G_αq/11_) mediate TGFβ activation in response to G-protein-coupled receptor (GPCR)-ligand binding as well as mechanical stretch in epithelial cells (Xu et al. 2009; John et al. 2016). GPCR signalling has also been implicated in normal alveologenesis (Funke et al. 2016). Cyclical mechanical stretch (CMS) has been shown to induce TGFβ activation in lung slices via a Rho-associated kinase (ROCK) - and αv integrin-dependent process (Froese et al. 2016), although the contribution to this by individual cell types is unknown. While stretch secondary to foetal breathing movements in utero is essential for early lung development (Donahoe, Longoni, and High 2016), the role of breathing-related CMS specifically in mesenchymal cells in alveolar development and the maintenance of adult alveoli has not been investigated.

We hypothesised that G_αq/11_ would mediate CMS-induced TGFβ activation via ROCK and integrin signalling in mesenchymal cells, and that this would be important in alveologenesis. Here we show, using mesenchymal G_αq/11_ knockout mouse models and an in vitro CMS system, that mesenchymal G_αq/11_ is essential for normal alveologenesis and maintenance of adult alveoli via CMS-induced TGFβ signalling, but that this occurs in a ROCK- and integrin-independent manner via a pathway likely to involve the TGFβ2 isoform.

## Results

### *Pdgfrb-Cre^+/−^;Gnaq^fl/fl^;Gna11^−/−^* mice are growth restricted and are not viable beyond P24

To understand whether mesenchymal G_αq/11_ deletion resulted in detrimental effects in vivo, gross phenotypes and genotype frequencies of offspring from the *Pdgfrb-Cre^+/−^ x Gnaq^fl/fl^;Gna11^−/−^* crosses were analysed. Fewer mesenchymal G knockout (*Pdgfrb-Cre^+/−^;Gnaq^fl/fl^;Gna11^−/−^*) pups reached genotyping age (P14) than was expected (6.6% observed compared with 12.5% expected, Chi squared value = 22.03, p<0.005, **Figure 1A**). Conversely, mice with at least one functional mesenchymal *Gnaq* or *Gna11* allele reached genotyping age at rates closer to the expected Mendelian frequencies (**Figure 1A**). Furthermore, *Pdgfrb-Cre^+/−^;Gnaq^fl/fl^;Gna11^−/−^* pups were notably smaller than littermates with at least one intact mesenchymal *Gnaq* or *Gna11* allele. *Pdgfrb-Cre^+/−^;Gnaq^fl/fl^;Gna11^−/−^* animals had a mean weight 1.9-3.2g lower than all other genotypes (5.4g vs 7.3-8.4g, p<0.03 **Figure 1B**). *Pdgfrb-Cre^+/−^;Gnaq^fl/fl^;Gna11^−/−^* pups were also smaller in physical size compared with control animals (**Figure 1C**). There was no sex-related difference in weight across genotypes (**Figure 1D**). These findings indicate that mesenchymal G_αq/11_ deletion causes a detrimental developmental phenotype, leading to death in utero or in early life.

**Figure 1:**
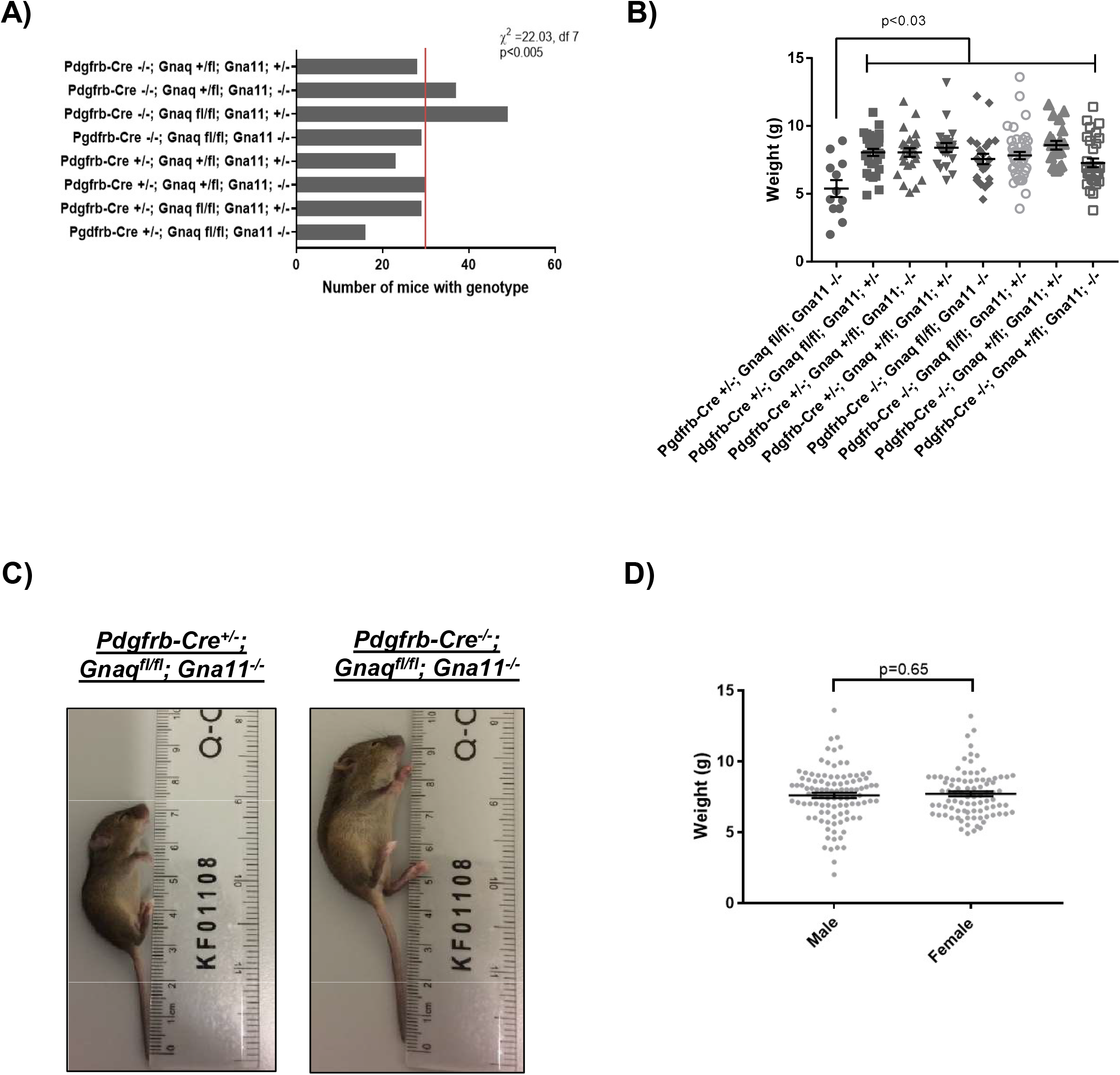
*Pdgfrb-Cre^+/−^;Gnaq^fl/fl^;Gna11^−/−^* mice are growth restricted A) Genotype frequencies from *Pdgfrb-Cre^+/−^* x *Gnaq^fl/fl^;Gna11^−/−^* breeding. Red line indicates the expected frequency for each genotype (n=30, 12.5%). Total n= 241, 24 litters, mean litter size 7.4. Chi-squared value (L^2^) = 22.03, degrees of freedom =7, p<0.005. B) Body weights of P14 pups by genotype. Mean ± SEM, one way ANOVA with Tukey’s multiple comparisons test, n = 12 *Pdgfrb-Cre^+/−^* x *Gnaq^fl/fl^;Gna11^−/−^* mice, n= 21-43 for other genotypes. C) Photograph of a P14 pup with the *Pdgfrb-Cre^+/−^; Gnaq^fl/fl^; Gna11^−/−^* genotype (left), and a *Gna11^−/−^* littermate (right). D) Body weights of all pups from *Pdgfrb-Cre / Gnaq^fl/fl^; Gna11^−/−^* crosses by sex at P14. Mean ± SEM, unpaired two-tailed Students T test, 88 female and 102 male mice.

The first two *Pdgfrb-Cre^+/−^;Gnaq^fl/fl^;Gna11^−/−^* mice from this breeding programme were humanely killed due to poor physical condition at P21 and P24. Therefore, all further analyses were performed in P14 mice, before evidence of ill health was observed. *Gnaq^fl/fl^;Gna11^−/−^* mice develop normally and do not express a phenotype (John et al. 2016), therefore *Gnaq^fl/fl^;Gna11^−/−^* littermates were used as controls for all analyses (from here referred to as *Gna11^−/−^* controls).

### *Pdgfrb-Cre^+/−^;Gnaq^fl/fl^;Gna11^−/−^* mice have impaired alveologenesis

To understand the role of mesenchymal G_αq/11_ signalling in lung development, the lungs of *Pdgfrb-Cre^+/−^;Gnaq^fl/fl^;Gna11^−/−^* mice and *Gna11^−/−^* controls were examined histologically. *Pdgfrb-Cre^+/−^;Gnaq^fl/fl^;Gna11^−/−^* mouse lungs exhibited clear abnormalities consistent with impaired alveolar development at P14 (**Figure 2A**). *Pdgfrb-Cre^+/−^;Gnaq^fl/fl^;Gna11^−/−^* lungs contained enlarged airspaces with a mean linear intercept distance of 63.47µm compared with 36.43µm in *Gna11^−/−^* mice (p=0.03, **Figure 2B**), thickened alveolar walls of 12.2µm compared with 7.0µm in *Gna11^−/−^* controls (p=0.03, **Figure 2C**), and fewer secondary crests (53.7 vs 107.2 per field, p=0.03, **Figure 2D**) relative to *Gna11^−/−^* littermate controls.

**Figure 2:**
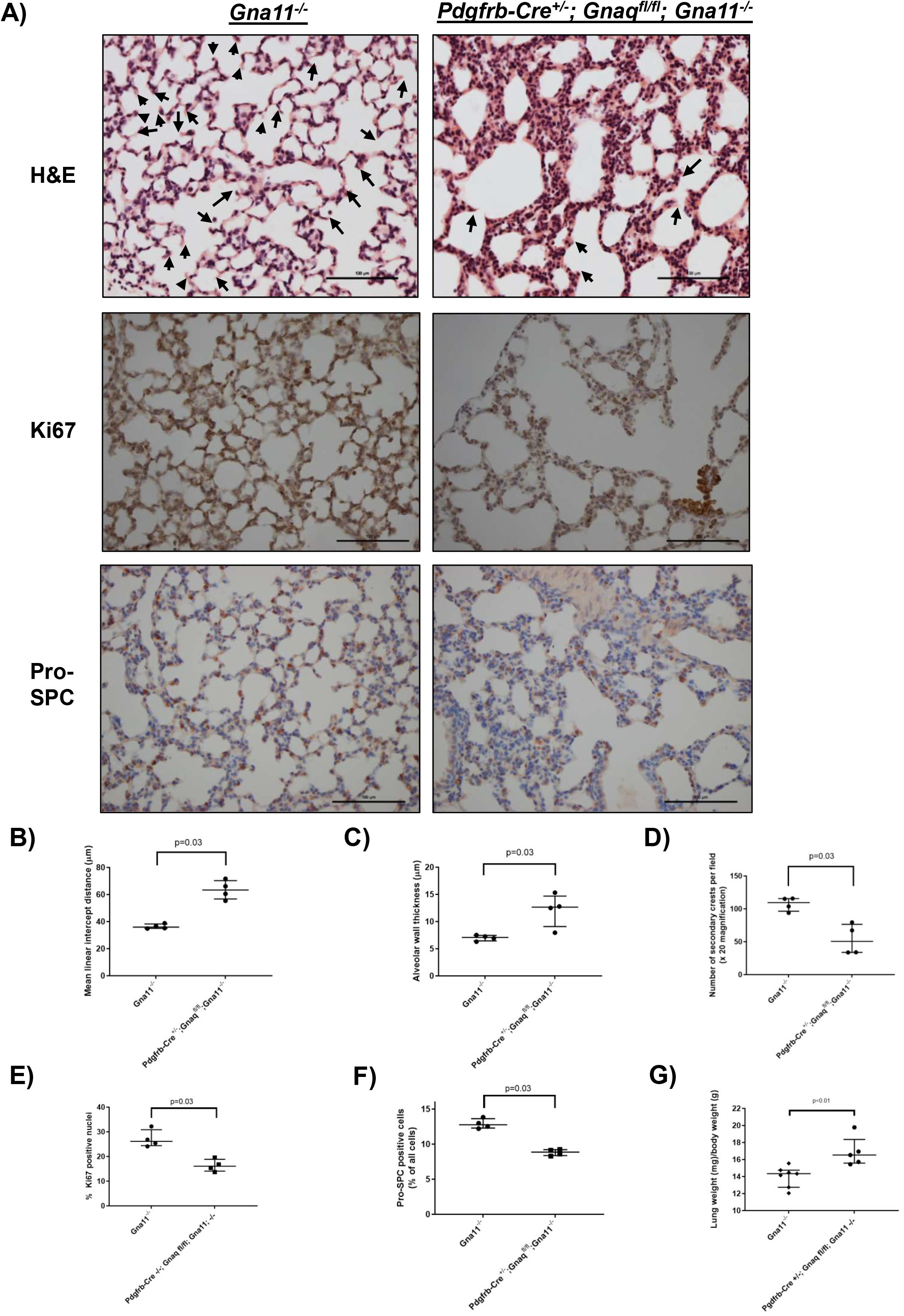
*Pdgfrb-Cre^+/−^;Gnaq^fl/fl^;Gna11^−/−^* mice have abnormal lung appearances characteristic of disturbed alveologenesis A) Haematoxylin and eosin (H&E) (top), Ki67 immunohistochemistry (middle), and pro-SPC immunohistochemistry (bottom) staining of lungs from a P14 *Gna11^−/−^* (left) and a *Pdgfrb-Cre^+/−^;Gnaq^fl/fl^;Gna11^−/−^* mouse (right). Arrows on H&E images indicate secondary crests. Images representative of 4 mice per group. Scale bars show 100µm. B) Mean linear intercept analysis of airspace size in P14 *Gna11^−/−^* and *Pdgfrb-Cre^+/−^;Gnaq^fl/fl^;Gna11^−/−^* mice. Median ± interquartile range, n=4 mice per group, two-tailed Mann Whitney test. C) Alveolar wall thickness in P14 *Gna11^−/−^* and *Pdgfrb-Cre^+/−^;Gnaq^fl/fl^;Gna11^−/−^* mice. Median ± interquartile range, n=4 mice per group, two-tailed Mann Whitney test. D) Quantification of the number of secondary crests per 20 x field in P14 *Gna11^−/−^* and *Pdgfrb-Cre^+/−^;Gnaq^fl/fl^;Gna11^−/−^* mice. Median ± interquartile range, n=4 mice per group, two-tailed Mann Whitney test. E) Quantification of Ki67 immunohistochemistry in P14 *Gna11^−/−^* and *Pdgfrb-Cre^+/−^;Gnaq^fl/fl^;Gna11^−/−^* mice. Shown as the percentage of Ki67 positive nuclei per 40x magnification field. Median ± interquartile range, n=4 mice per group, two-tailed Mann Whitney test. F) Quantification of Pro-SPC immunohistochemistry in P14 *Gna11^−/−^* and *Pdgfrb-Cre^+/−^;Gnaq^fl/fl^;Gna11^−/−^* mice. Shown as the percentage of pro-SPC positive cells per 40x magnification field. Median ± interquartile range, n=4 mice per group, two-tailed Mann Whitney test. G) Relative lung to body weights in P14 *Gna11^−/−^* and *Pdgfrb-Cre^+/−^;Gnaq^fl/fl^;Gna11^−/−^* mice. Median ± interquartile range, n=5 *Pdgfrb-Cre^+/−^;Gnaq^fl/fl^;Gna11^−/−^* mice, n=6 *Gna11^−/−^* controls, two-tailed Mann Whitney test.

In addition to these structural abnormalities, *Pdgfrb-Cre^+/−^;Gnaq^fl/fl^;Gna11^−/−^* lungs expressed lower levels of the proliferative marker Ki67 than *Gna11^−/−^* controls, with 16% of cell nuclei staining positively for Ki67 in *Pdgfrb-Cre^+/−^;Gnaq^fl/fl^;Gna11^−/−^* lungs compared with 26% in *Gna11^−/−^* controls (p=0.03, **Figure 2A, 2E**). Furthermore, *Pdgfrb-Cre^+/−^;Gnaq^fl/fl^;Gna11^−/−^* lungs contained a lower proportion of cells staining positively for the type II epithelial cell marker pro-surfactant protein C (pro-SPC) than *Gna11^−/−^* control lungs, at 8.9% and 12.8% of all cells, respectively (**Figure 2A, 2F**).

Finally, *Pdgfrb-Cre^+/−^;Gnaq^fl/fl^;Gna11^−/−^* lungs were heavier relative to total body weight compared with lungs from *Gna11^−/−^* mice (16.5 vs 14.3mg/g total body weight, p<0.01, **Figure 2G**), suggesting elevated lung density in these animals. Overall, these structural, proliferative, and cellular differentiation abnormalities indicate a disturbance to alveologenesis in *Pdgfrb-Cre^+/−^;Gnaq^fl/fl^;Gna11^−/−^* mice.

### Myofibroblast differentiation and function is defective in *Pdgfrb-Cre^+/−^;Gnaq^fl/fl^;Gna11^−/−^* mouse lungs

Myofibroblasts are essential for normal alveolar development, therefore studies were undertaken to assess myofibroblast differentiation and function in *Pdgfrb-Cre^+/−^;Gnaq^fl/fl^;Gna11^−/−^* lungs.

Immunohistochemical staining for the myofibroblast marker α-smooth muscle actin (αSMA) demonstrated fewer myofibroblasts in the lungs of P14 *Pdgfrb-Cre^+/−^;Gnaq^fl/fl^;Gna11^−/−^* mice compared with *Gna11^−/−^* littermate controls (**Figure 3A**). While overall αSMA staining was decreased in *Pdgfrb-Cre^+/−^;Gnaq^fl/fl^;Gna11^−/−^* lungs, there was no significant reduction in the proportion of αSMA-positive secondary crests compared with *Gna11^−/−^* lungs (0.69 vs 0.84 in controls, p =0.2, **Figure 3B**).

**Figure 3:**
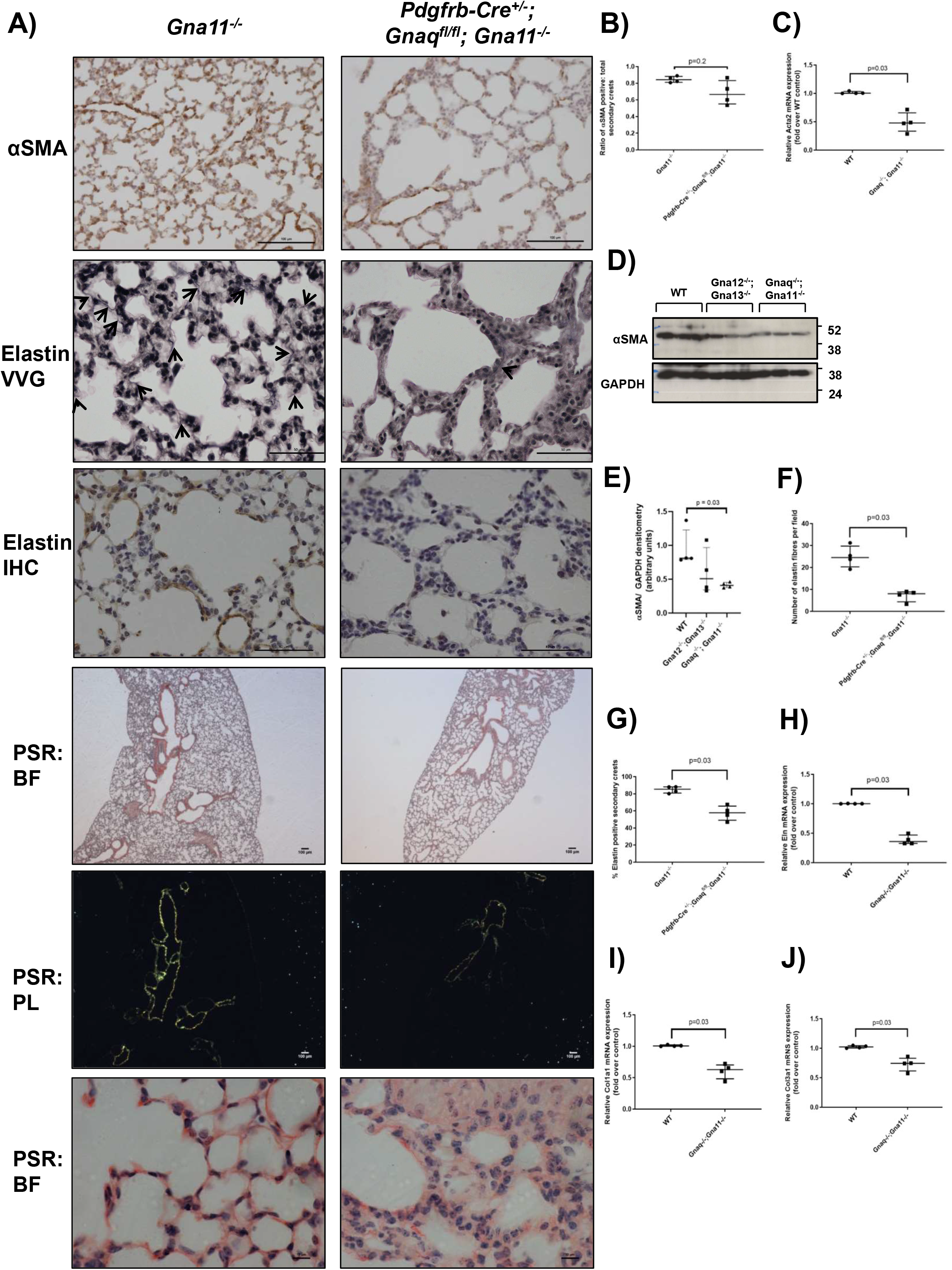
*Pdgfrb-Cre^+/−^;Gnaq^fl/fl^;Gna11^−/−^* mice have reduced lung myofibroblast differentiation and function A) αSMA immunohistochemistry (row 1), elastin Verhoeff van Gieson stain (row 2), elastin immunohistochemistry (row 3), and picrosirius red (PSR) staining (row 4-6) from P14 *Gna11^−/−^*(left) and *Pdgfrb-Cre^+/−^;Gnaq^fl/fl^;Gna11^−/−^*(right) mice. Arrows on elastin images shown elastin fibres. Picrosirius red images shown are bright field (BF, row 4), polarised light (row 5), and bright field at high magnification (row 5). Representative images from 4 mice per genotype. Scale bars show 100µm (αSMA, PSR), 50µm (elastin VVG and IHC), and 10µm (picrosirius red high magnification). B) Quantification of the proportion of secondary crests that stained positively for αSMA in P14 *Gna11^−/−^* and *Pdgfrb-Cre^+/−^;Gnaq^fl/fl^;Gna11^−/−^* lungs. Median ± interquartile range, n=4 mice per group, two-tailed Mann Whitney test. C) *Acta2* mRNA expression in WT, *Gna12^−/−^;Gna13^−/−^*, and *Gnaq^−/−^;Gna11^−/−^* MEFs. Median ± interquartile range, n=4 per group, two-tailed Mann Whitney test. D) Representative western blot showing αSMA expression in wild-type (WT), *Gna12^−/−^;Gna13^−/−^*, and *Gnaq^−/−^;Gna11^−/−^* MEFs. E) Densitometry of western blots of αSMA expression in wild-type (WT), *Gna12^−/−^;Gna13^−/−^*, and *Gnaq^−/−^;Gna11^−/−^* MEFs. Median ± interquartile range, n=4, two-tailed Mann Whitney test. F) The number of elastin fibres per high powered field (40 x magnification) in P14 *Gna11^−/−^* and *Pdgfrb-Cre^+/−^;Gnaq^fl/fl^;Gna11^−/−^* lungs. Median ± interquartile range, n=4 mice per group, two-tailed Mann Whitney test. G) The proportion of secondary crests that stained positively for elastin in each high powered field (40 x magnification) in P14 *Gna11^−/−^* and *Pdgfrb-Cre^+/−^;Gnaq^fl/fl^;Gna11^−/−^* lungs. Median ± interquartile range, n=4 mice per group, two-tailed Mann Whitney test. H) *Eln* mRNA expression in wild-type (WT) and *Gnaq^−/−^;Gna11^−/−^* MEFs. Median ± interquartile range, n=4, two-tailed Mann Whitney test. I) *Col1a1* mRNA expression in wild-type (WT) and *Gnaq^−/−^;Gna11^−/−^* MEFs. Median ± interquartile range, n=4, two-tailed Mann Whitney test. J) *Col3a1* mRNA expression in wild-type (WT) and *Gnaq^−/−^;Gna11^−/−^* MEFs. Median ± interquartile range, n=4, two-tailed Mann Whitney test. VVG = Verhoeff van Gieson; IHC = immunohistochemistry

To investigate whether G_αq/11_ knockout influences myofibroblast differentiation, murine embryonic fibroblasts (MEFs) that were wild-type (WT), G_αq/11_ deficient (*Gnaq^−/−^;Gna11^−/−^*) or G_α12/13_ deficient (*Gna12^−/−^;Gna13^−/−^*) were assessed for αSMA protein and Acta2 mRNA expression. MEFs with a long-term deficiency in G_αq/11_ had lower Acta2 mRNA (**Figure 3C**) and αSMA protein expression than WT MEFs, whereas MEFs lacking G_α12/13_, another G_α_ subunit family, did not have significantly different αSMA expression compared with WT cells (**Figure 3D, 3E**). This implies a key role for G_αq/11_ signalling in the differentiation of myofibroblasts from mesenchymal precursor cells.

*Pdgfrb-Cre^+/−^;Gnaq^fl/fl^;Gna11^−/−^* lungs also showed evidence of defective myofibroblast synthetic function. *Pdgfrb-Cre^+/−^;Gnaq^fl/fl^;Gna11^−/−^* lungs contained fewer elastin fibres (7.4 vs 24.9 fibres per field, p=0.03, **Figure 3A & 3F**) and fewer elastin-positive secondary crests (57.5% vs 84.8%, p=0.03, **Figure 3G**) than *Gna11^−/−^* mouse lungs. Furthermore, picrosirius red staining revealed that P14 *Pdgfrb-Cre^+/−^;Gnaq^fl/fl^;Gna11^−/−^* mouse lungs contained less collagen than the lungs of *Gna11^−/−^* controls (**Figure 3A**). These data were supported by lower *Eln*, *Col1a1* and *Col3a1* mRNA expression in *Gnaq^−/−^;Gna11^−/−^* MEFs than WT MEFs (**Figure 3H-J**). These findings imply a failure of myofibroblast differentiation in the lungs of mice lacking mesenchymal G_αq/11_ associated with a reduction in myofibroblast function, leading to a reduction in subepithelial matrix deposition.

### *Pdgfrb-Cre^+/−^;Gnaq^fl/fl^;Gna11^−/−^* mice have abnormal peripheral pulmonary vessels

Pericytes are important precursor cells to pulmonary myofibroblasts, and originate in the perivascular region. Therefore, we examined the pulmonary vasculature histologically to assess for abnormalities caused by mesenchymal G deletion. P14 *Pdgfrb-Cre^+/−^;Gnaq^fl/fl^;Gna11^−/−^* lungs contained markedly abnormal peripheral pulmonary vessels (**Figure 4A-G**), with significantly thicker walls than the peripheral pulmonary vessels of *Gna11^−/−^* controls (mean maximum wall thickness 16.4 vs 7.3µm, p=0.03, **Figure 4H**). These vessels consisted of a thin CD31 positive endothelial layer (**Figure 4B**) surrounded by a thickened αSMA positive vascular smooth muscle layer (**Figure 4C**) without increased proliferation (Ki67 positive; **Figure 4D**), indicating that the smooth muscle layer was hypertrophic rather than hyperplastic. These abnormal vessels did not contain significant collagen or elastin layers (**Figure 4E-G)**. In contrast, the alveolar capillaries of *Pdgfrb-Cre^+/−^;Gnaq^fl/fl^;Gna11^−/−^* lungs had a similar appearance to those seen in *Gna11^−/−^* lungs (**Figure 4J**).

**Figure 4:**
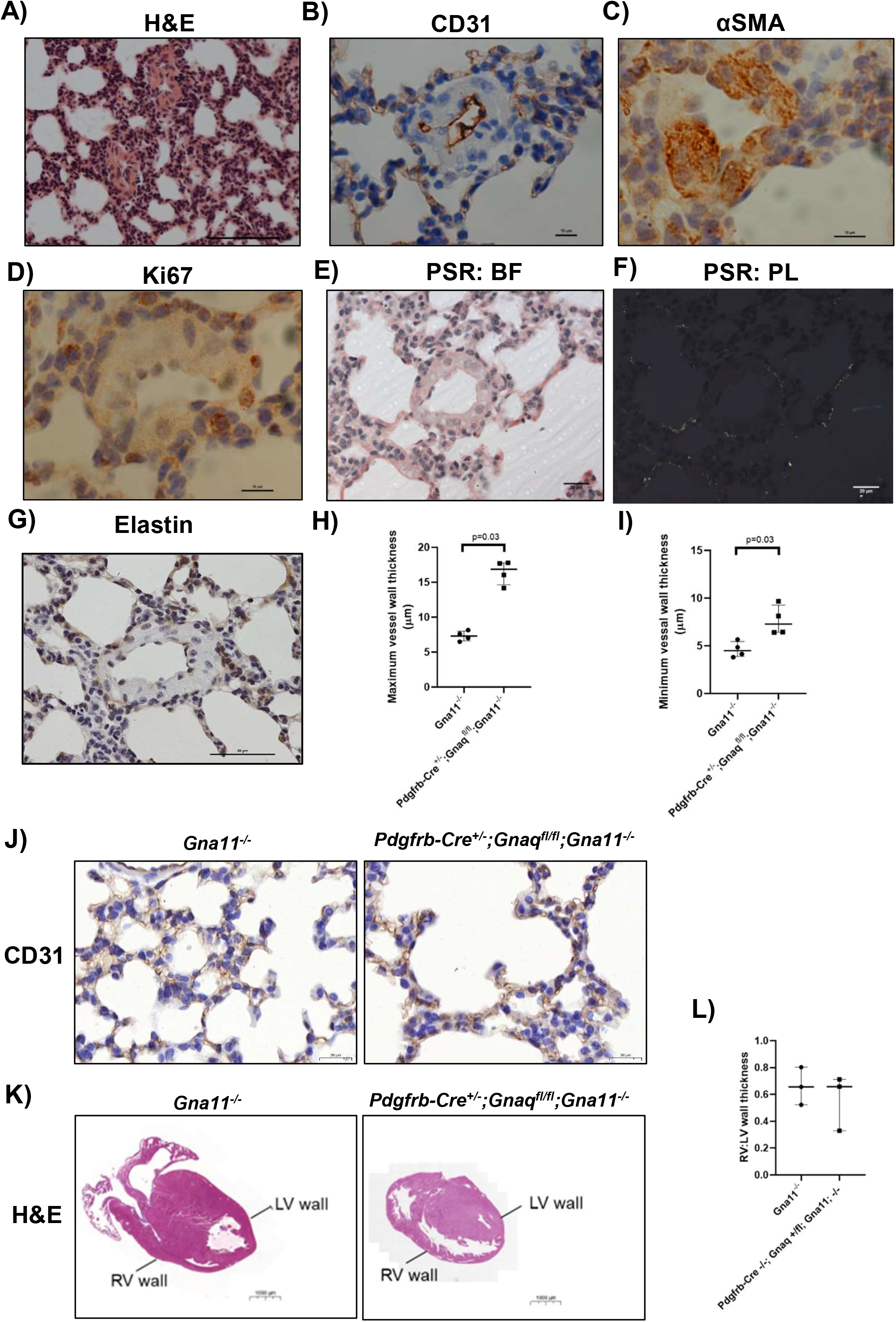
The lungs of *Pdgfrb-Cre^+/−^;Gnaq^fl/fl^;Gna11^−/−^* mice contain abnormal peripheral pulmonary vessels. A-G) Lung sections from P14 *Pdgfrb-Cre^+/−^;Gnaq^fl/fl^;Gna11^−/−^* mice were stained using various techniques. A) Haematoxylin and eosin stain. Scale bar shows 100µm. B) CD31 immunohistochemistry. Scale bar shows 10µm. C) αSMA immunohistochemistry. Scale bar shows 10µm. D) Ki67 immunohistochemistry. Scale bar shows 10µm. E) Picrosirius red stain (PSR). Same image shown using bright field (BF, E) and polarised light (PL, F) illumination). Scale bar shows 20µm. G) Elastin immunohistochemistry. Scale bar shows 50µm. H) Quantification of maximum peripheral vessel wall thickness in P14 *Gna11^−/−^* and *Pdgfrb-Cre^+/−^;Gnaq^fl/fl^;Gna11^−/−^* lungs. Median ± interquartile range, n=4 mice per group, two-tailed Mann Whitney test. I) Quantification of minimum peripheral vessel wall thickness in P14 *Gna11^−/−^* and *Pdgfrb-Cre^+/−^;Gnaq^fl/fl^;Gna11^−/−^* lungs. Median ± interquartile range, n=4 mice per group, two-tailed Mann Whitney test. J) CD31 immunohistochemistry from P14 *Gna11^−/−^* (left) and *Pdgfrb-Cre^+/−^;Gnaq^fl/fl^;Gna11^−/−^* (right) mice. Representative images from 4 mice per genotype. Scale bars show 20µm. K) Haematoxylin and eosin stain of representative hearts from P14 *Gna11^−/−^* (left) and *Pdgfrb-Cre^+/−^;Gnaq^fl/fl^;Gna11^−/−^* (right) mice. Scale bars show 1000µm. L) Right: left cardiac ventricular wall thickness ratios in P14 *Gna11^−/−^* (left) and *Pdgfrb-Cre^+/−^;Gnaq^fl/fl^;Gna11^−/−^* (right) mice. Median ± interquartile range, n=3 mice per group,

Given the similarity in appearance of the abnormal peripheral pulmonary vasculature in *Pdgfrb-Cre^+/−^;Gnaq^fl/fl^;Gna11^−/−^* lungs to those seen in pulmonary arterial hypertension, we assessed the hearts from these animals for evidence of right ventricular hypertrophy. We found no difference in right: left ventricular wall ratio in *Pdgfrb-Cre^+/−^;Gnaq^fl/fl^;Gna11^−/−^* mice relative to controls (**Figure 4K-L**). These data suggest a primary Pdgfrb^+^ cell-driven defect, rather than secondary pulmonary hypertension due to impaired alveologenesis.

### *Pdgfrb-Cre^+/−^;Gnaq^fl/fl^;Gna11^−/−^* mice have kidney abnormalities

As *Pdgfrb* expression is not exclusive to lung mesenchymal cells, the kidneys, hearts, livers, and bowel of *Pdgfrb-Cre^+/−^;Gnaq^fl/fl^;Gna11^−/−^* mice were assessed for extrapulmonary abnormalities.

We observed an expansion and prominence of medullary mesenchymal cells in *Pdgfrb-Cre^+/−^;Gnaq^fl/fl^;Gna11^−/−^* kidneys demonstrated by αSMA and PDGFRβ staining (**Figure 5A**), with associated thinning of the cortex (median cortex: medulla ratio 0.31 in *Pdgfrb-Cre^+/−^;Gnaq^fl/fl^;Gna11^−/−^* kidneys and 0.43 in *Gna11^−/−^* controls, p<0.03, **Figure 5B, C**). The relative kidney to total body weight values of *Pdgfrb-Cre^+/−^;Gnaq^fl/fl^;Gna11^−/−^* mouse kidneys were not different to *Gna11^−/−^* controls (median kidney: total body weight ratio 7.3 in *Pdgfrb-Cre^+/−^;Gnaq^fl/fl^;Gna11^−/−^* mice and 6.5 in *Gna11^−/−^* controls, p=0.55; **Figure 5D**). These data suggest that mesenchymal G_αq/11_ is important in normal kidney development.

**Figure 5:**
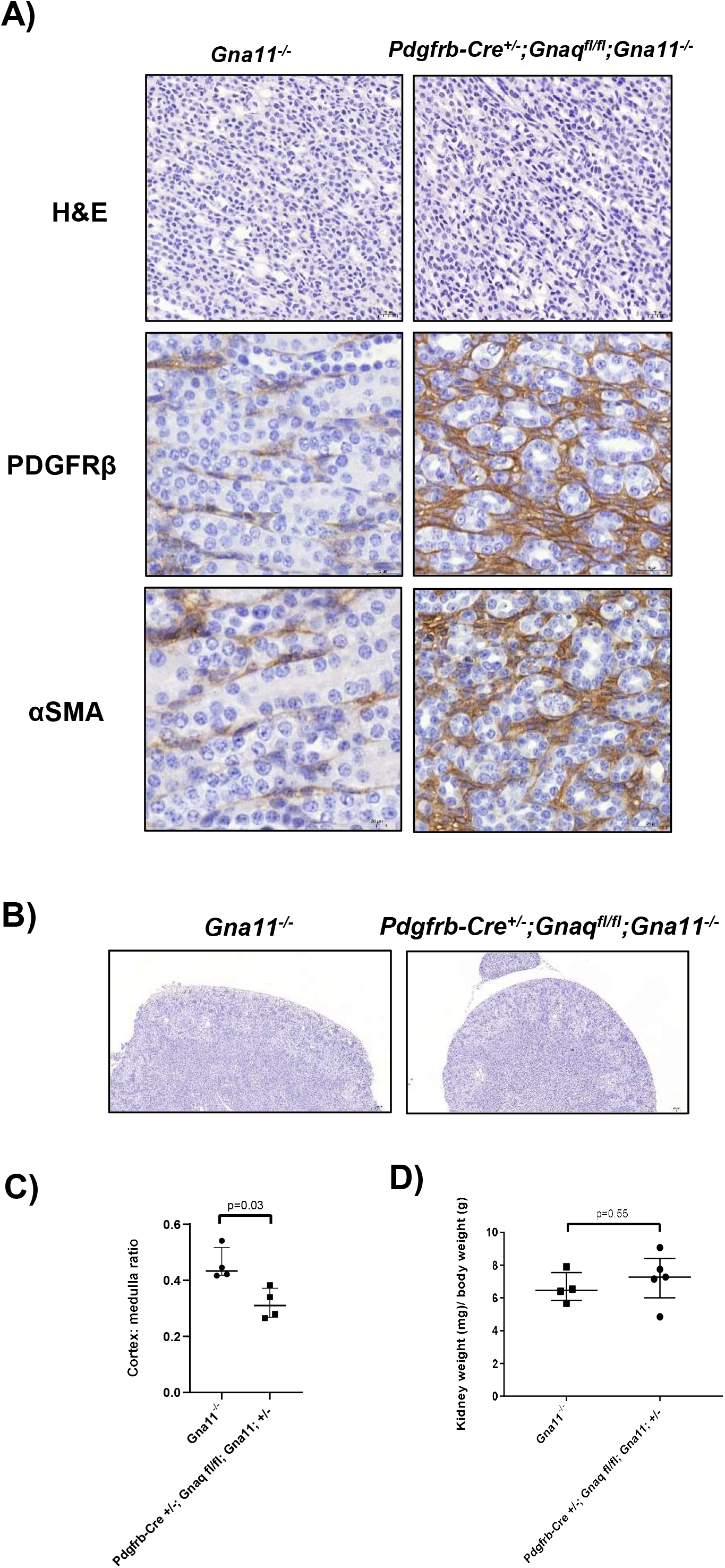
*Pdgfrb-Cre^+/−^;Gnaq^fl/fl^;Gna11^−/−^* have kidney abnormalities A) Haematoxylin and eosin, PDGFRβ immunohistochemistry, and αSMA immunohistochemistry of renal medulla, images of P14 *Gna11^−/−^* and *Pdgfrb-Cre^+/−^;Gnaq^fl/fl^;Gna11^-/^* mouse kidneys. Representative images from 4 mice per genotype. Scale bars show 20µm. B) Low magnification images of haematoxylin and eosin staining of P14 *Gna11^−/−^* (left) and *Pdgfrb-Cre^+/−^;Gnaq^fl/fl^;Gna11^−/−^* (right) mice. Scale bars show 200µm. C) Cortex: medulla ratios of P14 *Gna11^−/−^* and *Pdgfrb-Cre^+/−^;Gnaq^fl/fl^;Gna11^-/^* mice. Median ± interquartile range, n=4 mice per group, two-tailed Mann Whitney test. D) Relative kidney: total body weight in P14 *Gna11^−/−^* and *Pdgfrb-Cre^+/−^;Gnaq^fl/fl^;Gna11^-/^* mice.

*Pdgfrb-Cre^+/−^;Gnaq^fl/fl^;Gna11^−/−^* mice had normal heart, liver and bowel histology (Figure S1), suggesting that mesenchymal G_αq/11_ signalling is not required for normal heart, liver, or bowel development or homeostasis from conception to P14 in mice.

### Mice with mesenchymal G**_α_**_q/11_ knockout induced in adulthood have emphysema with altered ECM, but no extrapulmonary abnormalities

To assess whether the abnormalities seen in *Pdgfrb-Cre^+/−^;Gnaq^fl/fl^;Gna11^−/−^* mice were related solely to disturbed organ developmental processes or could also affect mature lungs, a tamoxifen-inducible mesenchymal G_αq/11_ knockout model (*Pdgfrb-Cre/ERT2^+/−^;Gnaq^fl/fl^;Gna11^−/−^*) was conducted in adult mice (**Figure 6A)**.

**Figure 6:**
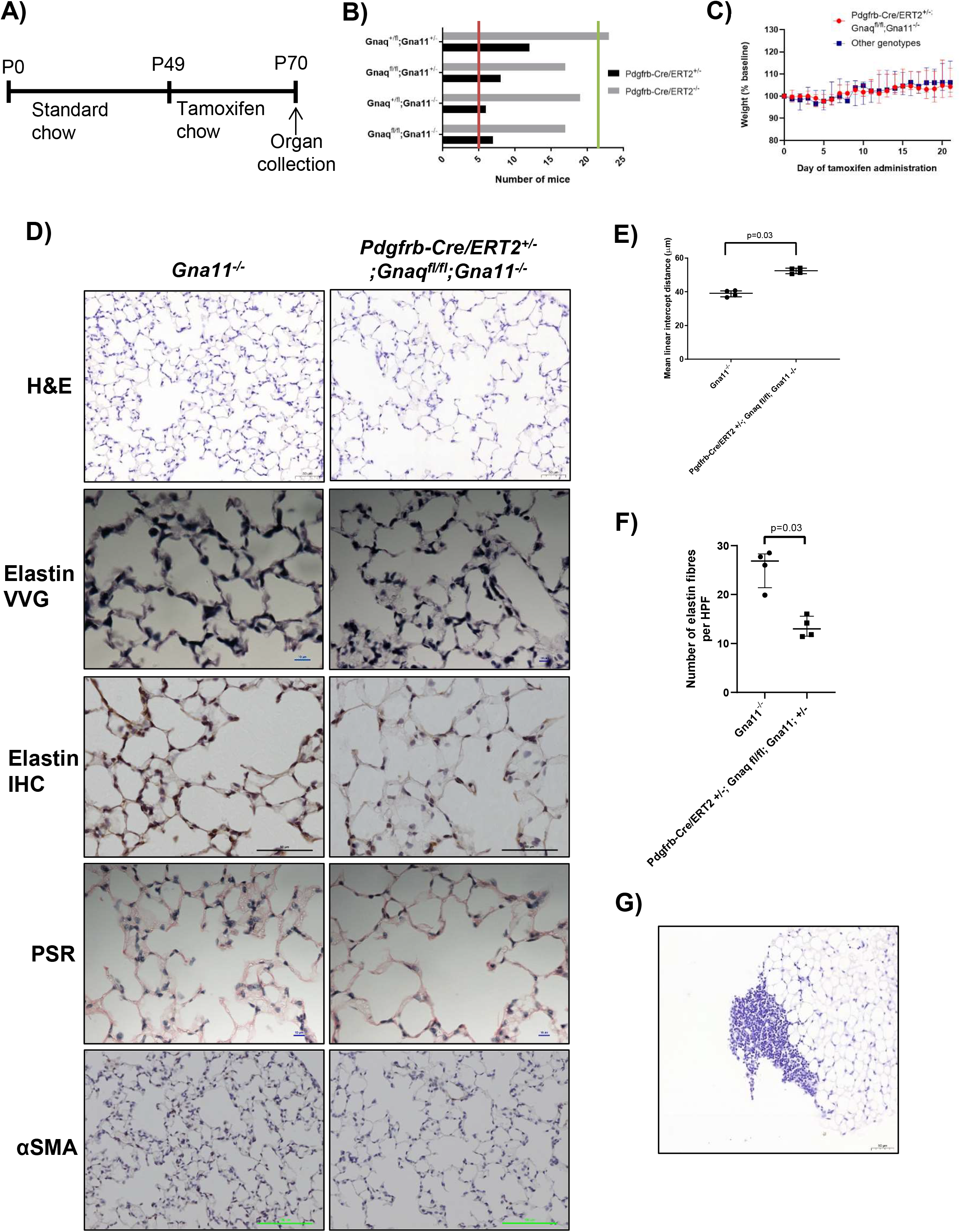
Mice with mesenchymal G_αq/11_ deletion in adulthood develop emphysema A) Protocol for tamoxifen administration in *Pdgfrb-Cre/ERT2^+/−^* x *Gnaq^fl/fl^;Gna11^−/−^* mouse colony. B) Genotype frequencies from *Pdgfrb-Cre/ERT2^+/−^* x *Gnaq^fl/fl^;Gna11^−/−^* breeding. Red line indicates the expected frequency of Pdgfrb-Cre/ERT2^+/−^ genotypes (5%, n =5), and green line indicates expected frequency of Pdgfrb-Cre/ERT2^−/−^ genotypes (20%, n=. 22, total n=109, 20 litters, mean litter size 5.5. C) Weights of *Pdgfrb-Cre/ERT2^+/−^;Gnaq^fl/fl^;Gna11^-/^* mice (red) and littermates of all other genotypes (blue) during 21 days of tamoxifen administration. D) Histology of lungs from *Gna11^−/−^* (left) and *Pdgfrb-Cre/ERT2^+/−^;Gnaq^fl/fl^;Gna11^-/^* (right) mice. Haematoxylin and eosin (row 1), elastin Verhoeff van Gieson (VVG, row 2), elastin immunohistochemistry (row 3), picrosirius red (PSR, row 4), and αSMA immunohistochemistry (row 5). Representative images from 4 mice per genotype. Scale bars show 50µm (H&E), 10µm (elastin VVG, PSR), 50µm (elastin IHC), and 100µm (αSMA IHC). E) Mean linear intercept distance in *Gna11^−/−^* and *Pdgfrb-Cre/ERT2^+/−^;Gnaq^fl/fl^;Gna11^-/^* mouse lungs. Median ± interquartile range, n=4 mice per group, two-tailed Mann Whitney test. F) Quantification of elastin fibres in *Gna11^−/−^* and *Pdgfrb-Cre/ERT2^+/−^;Gnaq^fl/fl^;Gna11^-/^* mouse lungs. Median ± interquartile range, n=4 mice per group, two-tailed Mann Whitney test. G) Representative image of mononuclear cell infiltrates seen in *Pdgfrb-Cre/ERT2^+/−^;Gnaq^fl/fl^;Gna11^-/^* mouse lungs.

Tamoxifen-naïve *Pdgfrb-Cre/ERT2^+/−^;Gnaq^fl/fl^;Gna11^−/−^* mice were born at the expected frequency. According to the supplier, it is expected that 20% of offspring from breeding of the Cre-expressing hemizygous mice with wild type mice will express the *Pdgfrb-Cre/ERT2* transgene (Laboratory), rather than the 50% Cre-expression rate observed in the germline *Pdgfrb-Cre^+/−^* mouse colony. The frequency of *Pdgfrb-Cre/ERT2^+/−^;Gnaq^fl/fl^;Gna11^−/−^* mice reaching genotyping age was 6.4%, compared with the expected 5% (total number of mice born 109; **Figure 6B)**. This indicates that having the *Pdgfrb-Cre/ERT2^+/−^;Gnaq^fl/fl^;Gna11^−/−^* genotype, without administration of tamoxifen, does not cause any gross developmental defects.

When a three week course of tamoxifen was administered to P49 *Pdgfrb-Cre/ERT2^+/−^;Gnaq^fl/fl^;Gna11^−/−^* mice (n=4, 1 female 3 male), no detrimental effect to health status was observed compared with littermate controls. Furthermore, *Pdgfrb-Cre/ERT2^+/^;Gnaq^fl/fl^;Gna11^−/−^* mice gained weight at the same rate as littermate controls with the other genotypes during the tamoxifen protocol (median weight on day 21 of tamoxifen 104.3% of baseline in *Pdgfrb-Cre/ERT2^+/−^;Gnaq^fl/fl^;Gna11^−/−^* mice compared to 106.2% of baseline in other genotypes, p=0.71; **Figure 6C**). A small reduction in weight was observed early in the tamoxifen protocol that was independent of genotype and was in keeping with a change in diet (Kiermayer et al. 2007). These data suggest that short-term mesenchymal G_αq/11_ knockout does not cause gross physiological disturbances *in vivo*.

On histological analysis, the lungs of *Pdgfrb-Cre/ERT2^+/−^;Gnaq^fl/fl^;Gna11^−/−^* mice treated with tamoxifen demonstrated increased airspace size compared with *Gna11^−/−^* controls (mean linear intercept distance 52.5µm in *Pdgfrb-Cre/ERT2^+/−^;Gnaq^fl/fl^;Gna11^−/−^* mice compared with 39.3µm in *Gna11^−/−^* controls, p=0.03, **Figure 6D, 6E**), suggestive of emphysema. *Pdgfrb-Cre/ERT2^+/−^;Gnaq^fl/fl^;Gna11^−/−^* lungs contained fewer elastin fibres than *Gna11^−/−^* controls after three weeks of tamoxifen (median number of elastin fibres per high powered field 13.0 in *Pdgfrb-Cre/ERT2^+/−^;Gnaq^fl/fl^;Gna11^−/−^* mice compared with 26.9 in *Gna11^−/−^* controls, p=0.03, **Figure 6D, 6F**), similar to the constitutive knockout. In contrast, *Pdgfrb-Cre/ERT2^+/−^;Gnaq^fl/fl^;Gna11^−/−^* lungs did not exhibit altered collagen deposition or evidence of fewer myofibroblasts (αSMA) when compared with *Gna11^−/−^* controls (**Figure 6D**). Three of the four *Pdgfrb-Cre/ERT2^+/−^;Gnaq^fl/fl^;Gna11^−/−^* mice also exhibited abnormal pulmonary mononuclear cellular aggregates which predominated at the pleural surfaces (**Figure 6G**), and were not observed in littermate control mice. Despite these abnormalities, *Pdgfrb-Cre/ERT2^+/−^;Gnaq^fl/fl^;Gna11^−/−^* mice did not exhibit signs of respiratory distress.

In contrast with *Pdgfrb-Cre^+/−^;Gnaq^fl/fl^;Gna11^−/−^* mice, *Pdgfrb-Cre/ERT2^+/−^;Gnaq^fl/fl^;Gna11^−/−^* mice administered tamoxifen did not exhibit any renal abnormalities on histology (**Figure S2**). This implies that mesenchymal G_αq/11_ is needed for normal kidney development, but not maintenance of the normal kidney.

### Cyclical mechanical stretch-induced TGF**β** activation in fibroblasts requires G**_α_**_q/11,_ but not ROCK or **α**v or **β**1 integrins

Given the crucial roles TGFβ in alveolar development, lung repair, and pericyte migration and differentiation, we investigated the role of mesenchymal G_αq/11_ in a cyclical stretch model of TGFβ activation. Mesenchymal cells with and without intact G_αq/11_ signalling were subjected to breathing-related CMS and TGFβ signalling was assessed. CMS-induced TGFβ signalling, as assessed by Smad2 phosphorylation, was significantly reduced in *Gnaq^−/−^;Gna11^−/−^* MEFs compared with WT MEFs (**Figure 7A-B)**. This finding was specific to the G_αq/11_ family of G proteins, as there was no effect of G_α12/13_ knockdown on stretch-induced TGFβ signalling in MEFs (**Figure 7A**).

**Figure 7:**
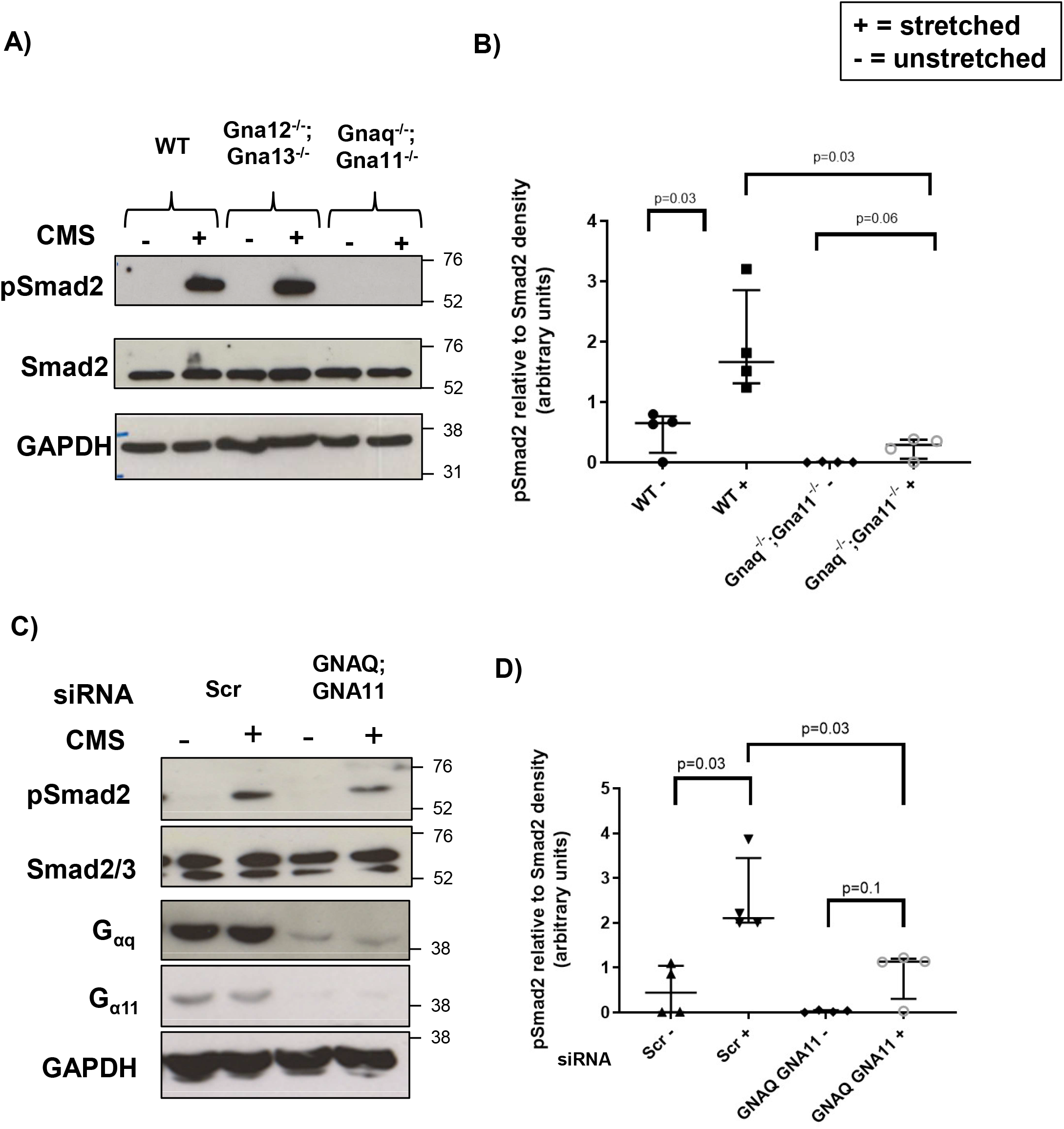
G_αq/11_ mediates stretch-induced TGFβ signalling in murine and human fibroblasts A) Representative western blot showing pSmad2 expression in wild-type (WT), *Gna12^−/−^;Gna13^−/−^*, and *Gnaq^−/−^Gna11^−/−^* MEFs subject to cyclical stretch (15% elongation, 1Hz, 48 hours). B) Densitometry of western blots from stretched MEFs shown as pSmad2 relative to Smad2 expression from 4 independent experiments. Median ± interquartile range, n=4, two-tailed Mann Whitney Test. C) Representative western blot showing pSmad2 expression in HLFs treated with non-targeting (Scr) or *GNAQ* and *GNA11* siRNA then subject to cyclical stretch (15% elongation, 0.3Hz, 24 hours). D) Densitometry of western blots from stretched HLFs shown as pSmad2 relative to Smad2 expression from 4 independent experiments. Median ± interquartile range, n=4, two-tailed Mann Whitney Test. + = stretched; - = unstretched

To validate the role of G_αq/11_ in stretch-induced TGFβ signalling in mesenchymal cells across species, human lung fibroblasts (HLFs) with and without siRNA-induced *GNAQ* and *GNA11* knockdown were subjected to breathing-related CMS. *GNAQ* and *GNA11* siRNA led to substantial reductions in both G_αq_ and G_α11_ protein expression in HLFs, and significantly reduced CMS-induced TGFβ signalling compared with scrambled control (Scr) siRNA as measured by phosphorylation of Smad2 (**Figure 7C-D**). These data indicate that G_αq/11_ is a key component of CMS-induced TGFβ signalling in both murine and human fibroblasts.

Previous studies have reported that G_αq/11_-induces TGFβ activation via the Rho-ROCK cascade and αv integrins in epithelial cells (Xu et al. 2009; Froese et al. 2016). As αvβ1, αvβ3, and αvβ5 integrins are expressed by myofibroblasts and are involved in TGFβ activation (Pakshir et al. 2020), we utilised chemical inhibition of these integrins and ROCK in our CMS model. When human fibroblasts were subject to breathing-related CMS in the presence of a ROCK1/2 inhibitor (Y27632), a pan αv integrin inhibitor (CWHM-12) or a β1 integrin-specific inhibitor (NOTT199SS), CMS-induced TGFβ signalling was not reduced (**Figure S3**). These data imply a novel pathway for CMS-induced TGFβ signalling in mesenchymal cells which requires G_αq/11_, but is independent of ROCK and integrin signalling.

### G**_α_**_q/11_ induces TGF**β**2 production, which is then available for CMS-induced serine protease-mediated activation

Proteases can activate latent TGFβ independently of integrins, therefore we assessed the effect of protease inhibitors in our CMS-induced TGFβ signalling system. A pan serine protease inhibitor 4-(2-aminoethyl)benzenesulfonyl fluoride (AEBSF), decreased CMS-induced Smad2 phosphorylation in HLFs (**Figure 8A-B**), whereas the MMP inhibitor GM-6001 had no effect on CMS-induced TGFβ signalling even at high concentrations (**Figure 8C-D**). These findings indicate that serine proteases mediate CMS-induced TGFβ signalling in mesenchymal cells.

**Figure 8:**
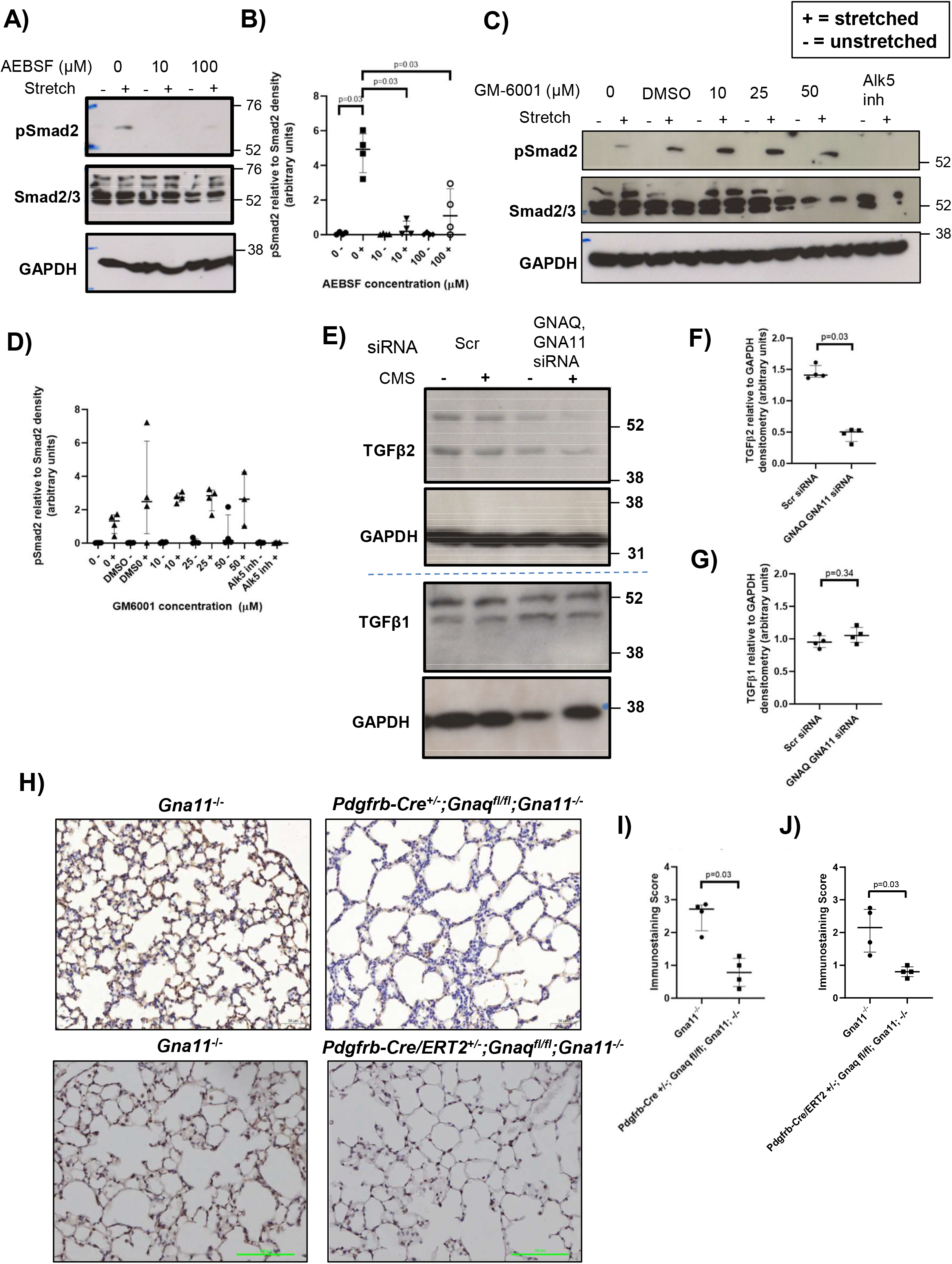
G_αq/11_ signalling induces the production of TGFβ2 which is then available for stretch-induced serine protease-mediated activation A) Representative pSmad2 western blot of human lung fibroblasts treated with the serine protease inhibitor AEBSF then subject to cyclical stretch (15% elongation, 0.3Hz, 48 hours). B) Relative pSmad2 to Smad2 densitometry of human lung fibroblasts treated with AEBSF then subject to cyclical stretch. Median ± interquartile range, n=4, two-tailed Mann Whitney test. C) Representative pSmad2 western blot of human lung fibroblasts treated with the matrix metalloproteinase inhibitor GM6001 then subject to cyclical stretch (15% elongation, 0.3Hz, 48 hours). D) Relative pSmad2 to Smad2 densitometry from human lung fibroblasts treated with GM6001 then subject to cyclical stretch. Median ± interquartile range, n=4, two-tailed Mann Whitney test. E) Representative TGFβ2 (top) and TGFβ1 (bottom) western blots of human lung fibroblasts subject to non-targeting (Scr) or *GNAQ* and *GNA11* siRNA and cyclical stretch (15% elongation, 0.3Hz, 24 hours). F) Relative TGFβ2 to GAPDH densitometry of human lung fibroblasts with and without siRNA-induced *GNAQ* and *GNA11* knockdown. Median ± interquartile range, n=4, two-tailed Mann Whitney test G) Relative TGFβ1 to GAPDH densitometry of human lung fibroblasts with and without siRNA-induced *GNAQ* and *GNA11* knockdown. Median ± interquartile range, n=4, two-tailed Mann Whitney test. H) TGFβ2 immunohistochemistry on P14 *Gna11^−/−^* (left) and *Pdgfrb-Cre^+/−^;Gnaq^fl/fl^;Gna11^−/−^* (right) mouse lungs (top row), and tamoxifen-treated P70 *Gna11^−/−^* (left) and *Pdgfrb-Cre/ERT2^+/−^;Gnaq^fl/fl^;Gna11^−/−^* mouse lungs. I) TGFβ2 immunohistochemistry scores of P14 *Gna11^−/−^* (left) and *Pdgfrb-Cre^+/−^;Gnaq^fl/fl^;Gna11^−/−^* (right) mouse lungs. Median ± interquartile range, n=4, two-tailed Mann Whitney test. J) TGFβ2 immunohistochemistry scores of tamoxifen-treated P70 *Gna11^−/−^* (left) and *Pdgfrb-Cre/ERT2^+/−^;Gnaq^fl/fl^;Gna11^−/−^*(right) mouse lungs. Median ± interquartile range, n=4, two-tailed Mann Whitney test. + = stretched; - = unstretched.

As TGFβ2 is the only TGFβ isoform that is not activated by integrins (Jenkins 2008), we hypothesised that breathing-related CMS would predominantly activate the TGFβ2 isoform in mesenchymal cells. While CMS did not influence TGFβ2 protein expression in HLFs, HLFs with siRNA-induced *GNAQ* and *GNA11* knockdown expressed less TGFβ2 than HLFs with intact G_αq/11_ signalling (**Figure 8E-F**), suggesting that G_αq/11_ plays a role in TGFβ2 production. Conversely, TGFβ1 protein expression was not affected by *GNAQ* and *GNA11* knockdown in HLFs (**Figure 8G**), suggesting an isoform-specific effect.

To evaluate the role of this CMS-induced TGFβ2 signalling pathway in alveologenesis, we assessed TGFβ2 expression in the lungs of mice from our mouse models. *Pdgfrb-Cre^+/−^;Gnaq^fl/fl^;Gna11^−/−^* lungs had a significantly lower TGFβ2 immunostaining score than *Gna11^−/−^* control lungs (median immunostaining score 0.8 in *Pdgfrb-Cre/ERT2^+/−^;Gnaq^fl/fl^;Gna11^−/−^* lungs, compared with 2.7 in *Gna11^−/−^* controls, p<0.03, **Figure 8I**). Similarly, *Pdgfrb-Cre/ERT2^+/−^;Gnaq^fl/fl^;Gna11^−/−^* mouse lungs also had reduced TGFβ2 deposition compared with Gna11^−/−^ controls after 3 weeks of tamoxifen (median immunostaining score 0.8 in *Pdgfrb-Cre/ERT2^+/−^;Gnaq^fl/fl^;Gna11^−/−^* lungs compared with 2.2 in *Gna11^−/−^* controls, p<0.03, **Figure 8J**). These data demonstrate that lungs lacking mesenchymal G_αq/11_ have less TGFβ2 available for breathing-related CMS-induced activation, and this may be important in alveologenesis and the maintenance of normal lung structure in vivo.

### G**_α_**_q/11_ influences expression of PDGF signalling components

Platelet-derived growth factor (PDGF) signalling is known to be important in alveolar development, and interacts with TGFβ signalling in normal development and disease (Gouveia, Betsholtz, and Andrae 2017, 2018). We therefore investigated how G_αq/11_ signalling influences the expression of PDGF signalling components in fibroblasts.

*Gnaq^−/−^;Gna11^−/−^* MEFs expressed significantly lower levels of *Pdgfb* and *Pdgfd* mRNA compared with wild-type cells (p=0.03, **Figure 9B, 9D**). There was not a statistically significant difference in the expression of *Pdgfa*, *Pdgfc*, *Pdgfra*, or *Pdgfrb* mRNA expression between *Gnaq^−/−^;Gna11^−/−^* and wild-type MEFs (**Figure 9A,C,E,F**), although there was a trend to reduced *Pdgfa* expression in *Gnaq^−/−^;Gna11^−/−^* MEFs (p=0.06, **Figure 9A**). These data imply that mesenchymal G_αq/11_ deletion influences the expression of PDGF signalling components, and thus may regulate PDGF signalling.

**Figure 9:**
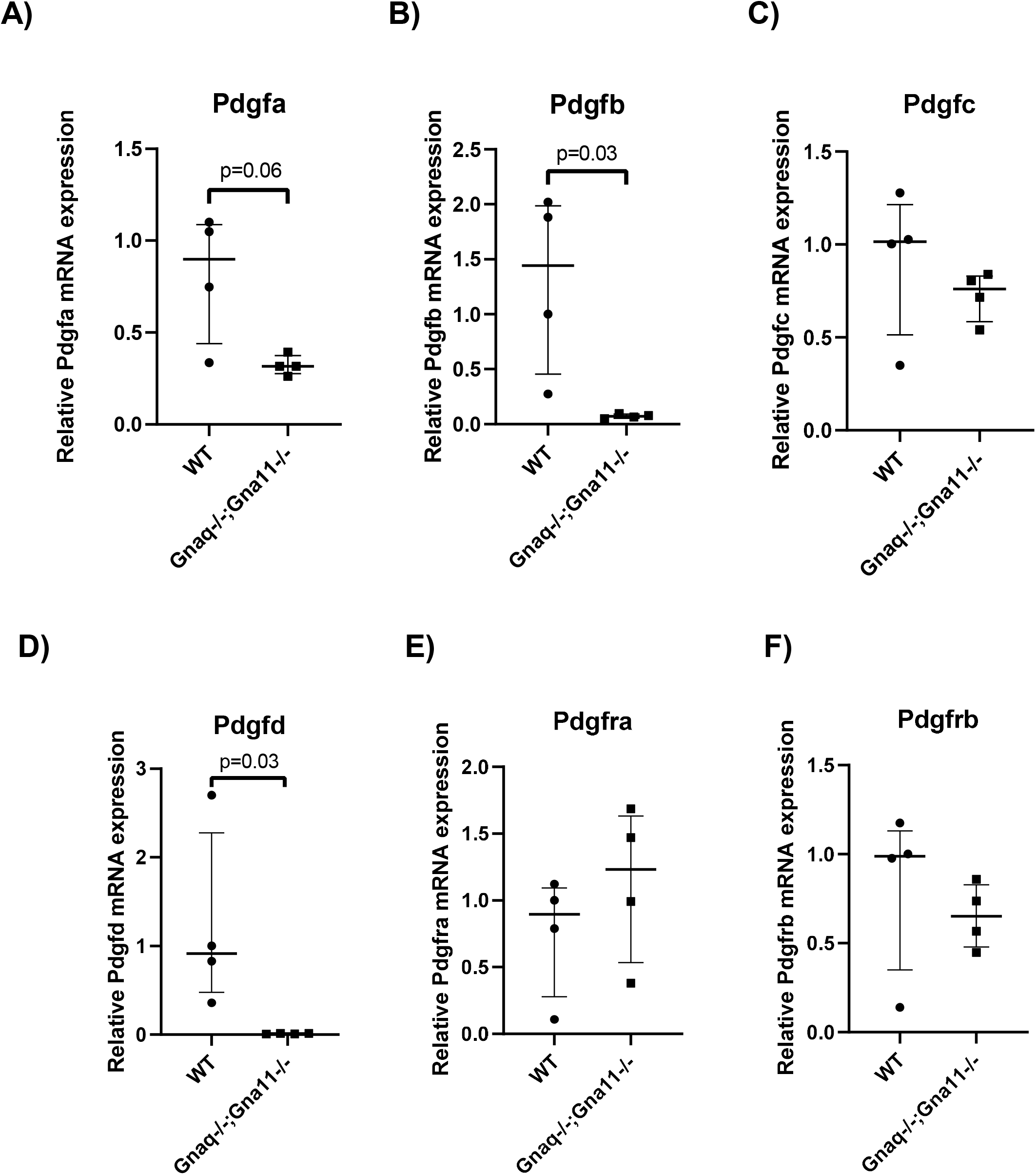
G_αq/11_ deletion influences expression of some PDGF transcripts in MEFs Relative mRNA expression of Pdgfa (A), Pdgfb (B), Pdgfc (C), Pdgfd (D), Pgdfra (E), and Pdgfrb (F) in wild-type (WT) and Gnaq^−/−^:Gna11^+/−^ MEFs. Median ± interquartile range, n=4, two-tailed Mann Whitney test.

## Discussion

In this study, we used mice with a targeted deletion of G_αq/11_ in mesenchymal cells to demonstrate that mesenchymal G_αq/11_ is essential for the development and maintenance of normal alveoli. Loss of G_αq/11_-mediated signalling in mesenchymal cells caused failure of the myofibroblast differentiation and ECM synthetic function required for alveolar development and the maintenance of the adult lung, and reduced mesenchymal cell TGFβ2 production is a key factor in these processes. In the absence of mesenchymal G_αq/11,_ TGFβ2 is unavailable for activation by CMS-induced serine proteases thereby diminishing downstream TGFβ signalling in both developing and adult lungs. These findings establish a previously undescribed role for breathing-related CMS in TGFβ2 generation, and suggest a role for TGFβ2 in alveolar development and lung homeostasis.

The role of G_αq/11_ in alveolar development has not previously been investigated, primarily because germline G_αq/11_ deletion is embryonically lethal (Offermanns et al. 1998) and murine alveolarisation occurs entirely postnatally (Beauchemin et al. 2016). Cell type-specific *Gnaq* and *Gna11* deletion in neural, cardiovascular, and haematological tissues have various manifestations ranging from no phenotype to profound cardiac abnormalities associated with perinatal death (Wettschureck et al. 2006; Wettschureck et al. 2004; Hoyer et al. 2010; Sassmann et al. 2010; Wettschureck et al. 2005; Wettschureck et al. 2007; Wettschureck et al. 2001). However, alveolar abnormalities have not been described in germline or conditional G_αq/11_ knockout mice, suggesting a unique role for mesenchymal G_αq/11_ in alveolar development and maintenance.

We propose that the key mechanisms underlying the abnormal alveologenesis and emphysema in mice with mesenchymal G_αq/11_ deletion present from conception or induced in adulthood, respectively, are failure of myofibroblast differentiation and synthetic function. Both *Pdgfrb-Cre^+/−^;Gnaq^fl/fl^;Gna11^−/−^* and *Pdgfrb-Cre/ERT2^+/−^;Gnaq^fl/fl^;Gna11^−/−^* mice had lower lung elastin deposition that controls. *Pdgfrb-Cre^+/−^;Gnaq^fl/fl^;Gna11^−/−^* lungs also contained fewer myofibroblasts and less collagen compared with controls, and mesenchymal cells lacking G_αq/11_ express less *Col1a1*, *Col3a1*, and *Eln* mRNA than cells with intact G_αq/11_. As myofibroblasts induce secondary septation by depositing ECM proteins at the tips of developing secondary septae, and loss of elastin is a key feature of emphysema (Ito et al. 2019), these data suggest that mesenchymal G_αq/11_-induced myofibroblast differentiation and function are required for alveolar development and homeostasis.

Secondary crest myofibroblasts (SCMFs) are known to derive from PDGFRα-expressing precursors (Boström et al. 1996; Lindahl et al. 1997; McGowan et al. 2008; R. Li et al. 2018), however the role of PDGRFβ^+^precursors in the development of SCMFs has not been described. While this study cannot definitively conclude that PDGFRβ^+^ precursors, such as pericytes, differentiate into SCMFs, it does show a role for PDGFRβ^+^ cells in alveolarisation. Whether this occurs via direct differentiation of SCMFs from PDGFRβ^+^ precursors, or via paracrine signalling from PDGFRβ^+^ cells should be the topic of further study.

*Pdgfrb-Cre^+/−^;Gnaq^fl/fl^;Gna11^−/−^* mouse lungs also contained abnormal peripheral pulmonary vessels, with a hypertrophic vascular smooth muscle layer. This could be explained by hypoxaemia-induced pulmonary arterial hypertension (PAH) secondary to the profound pulmonary defects, in combination with disturbed GPCR signalling, resulting in vascular remodelling (Patel et al. 2018; Cheng et al. 2012). However, *Pdgfrb-Cre^+/−^;Gnaq^fl/fl^;Gna11^−/−^* mice did not exhibit signs of respiratory distress at P14, and cardiac histology did not show evidence of right ventricular hypertrophy, which would be expected in PAH. We hypothesise that G_αq/11_ deletion prevents pericytes from migrating away from the perivascular region to the alveolar parenchyma, resulting in dysregulated vascular smooth muscle growth. However, firm conclusions on the cause of the abnormal peripheral pulmonary vessels in *Pdgfrb-Cre^+/−^;Gnaq^fl/fl^;Gna11^−/−^* mice cannot be drawn from this work.

Altered CMS-induced TGFβ activation is likely to be a key driver of the lung phenotypes observed in *Pdgfrb-Cre^+/−^;Gnaq^fl/fl^;Gna11^−/−^* and *Pdgfrb-Cre^+/−^;Gnaq^fl/fl^;Gna11^−/−^* mice. TGFβ drives myofibroblast differentiation, cellular migration, and ECM protein production (Harrell et al. 2018), and deficiencies and genetic polymorphisms in TGFβ signalling pathway components have been associated with emphysema (Bonniaud et al. 2004; M. Li et al. 2011; Hersh et al. 2009; Celedón et al. 2004). Both lung stretch and tightly-controlled TGFβ signalling are important for normal lung development and regeneration (Belcastro et al. 2015; Nakanishi et al. 2007; Chen et al. 2005; Chen et al. 2008; Sterner-Kock et al. 2002; Pieretti et al. 2014; Deng et al. 2019; Gauldie et al. 2003; Vicencio et al. 2004; Alejandre-Alcázar et al. 2008; Bonniaud et al. 2004; Donahoe, Longoni, and High 2016), and CMS has been demonstrated to induce TGFβ signalling in a number of models and organ systems (Froese et al. 2016; John et al. 2016; Fujita et al. 2010; Furumatsu et al. 2013; Maeda et al. 2011; Russo et al. 2018; Wang et al. 2013). Using the same *Gnaq^fl/fl^;Gna11^−/−^* mice used in our study, John et al described age-related emphysema related to reduced stretch-induced TGFβ signalling in mice lacking G_αq/11_ in type II alveolar epithelial cells (John et al. 2016). Open access RNA-Seq data on the LungMAP and IPF Cell Atlas databases show that in human and mouse lung, PDGFRβ-positive cells include pericytes, fibroblasts and myofibroblasts (www.ipfcellatlas.com; www.lungmap.net). We therefore used human lung fibroblasts and murine embryonic fibroblasts to assess the role of mesenchymal G_αq/11_ in CMS-induced TGFβ signalling, and to demonstrate the generalisability of our findings across species.

CMS-induced TGFβ signalling in mesenchymal cells was dependent on serine proteases and independent of αv integrins, contrary to previous work in lung slices and epithelial cells (Froese et al. 2016; Xu et al. 2009). This indicated that TGFβ2, an isoform that is activated by proteases but not integrins (Jenkins 2008), may be the primary TGFβ isoform activated by mesenchymal cell stretch. G_αq/11_-deficient fibroblasts expressed less TGFβ2, but had unchanged levels of TGFβ1, compared with cells that express G_αq/11_, suggesting a TGFβ isoform-specific effect of G_αq/11_ deletion. These data suggest a novel pathway in which mesenchymal G_αq/11_ drives TGFβ2 production, which is then available for protease-mediated activation.

This is the first study to propose an isoform-specific role for TGFβ2 in mammalian alveolar development and lung homeostasis. The three TGFβ isoforms are highly expressed during lung development with distinct spatial and temporal expression patterns (Schmid et al. 1991), however little is known about the specific regulation of TGFβ2 signalling. *Tgfb2^−/−^* mice die shortly after birth from developmental abnormalities distinct from those seen in *Tgfb1^−/−^* or *Tgfb3^−/−^* mice (Sanford et al. 1997; Shull et al. 1992; Kaartinen et al. 1995). *Tgfb2^−/−^* mice have no gross lung morphological abnormalities in late intrauterine gestation, however collapsed conducting airways are found postnatally (Sanford et al. 1997). While the *Pdgfrb-Cre^+/−^;Gnaq^fl/fl^;Gna11^−/−^* mice generated in the present study did not share phenotypic features with*Tgfb2^−/−^* mice, it is possible that TGFβ2 production by non-mesenchymal cells is sufficient for normal prenatal development. Additionally, as alveolarisation occurs entirely postnatally in mice, the role of TGFβ2 in alveolar development that we descrube could not be observed in *Tgfb2^−/−^* mice due to perinatal death. Our data demonstrate that loss of mesenchymal G_αq/11_ causes a loss of the precise control of TGFβ signalling in the lungs, resulting in abnormal alveologenesis and loss of lung homeostasis in developed lungs. Further work is required to understand the precise roles of individual TGFβ isoforms in these processes.

The PDGF family is known be important in lung development and regeneration, with PDGFA being particularly important in alveolar development (Gouveia, Betsholtz, and Andrae 2018; Gouveia et al. 2020; Gokey et al. 2021). We found a trend towards reduced *Pdgfa* expression in MEFs with G_αq/11_ deletion, as well as *Pdgfb* and *Pdgfc*, suggesting that G_αq/11_ signalling may interact with PDGF-related pathways. Postnatal deletion of *Pdgfra*, which encodes the major receptor for PDGFA, reduces lung *Tgfb2*, but not *Tgfb1,* transcripts (C. Li et al. 2019), further supporting a role for PDGF signalling in G_αq/11_- and TGFβ2-driven alveolar development and regeneration. However, elastin deposition during alveologenesis may not be dependent on PDGFA (Gouveia et al. 2020), therefore PDGF-independent pathways are also likely to be involved in driving the abnormalities in *Pdgfrb-Cre^+/−^;Gnaq^fl/fl^;Gna11^−/−^* and *Pdgfrb-Cre/ERT2^+/−^;Gnaq^fl/fl^;Gna11^−/−^* mouse lungs. As pulmonary mesenchymal cells are predominantly PDGF receptor-expressing, rather than PDGF ligand producing (Gouveia, Betsholtz, and Andrae 2017), and G_αq/11_ deletion did not alter *Pdgfra* or *Pdgfrb* expression, we hypothesise that mesenchymal G_αq/11_ deletion reduces lung TGFβ2 signalling, which subsequently alters PDGF ligand expression by other cell types. However, it was beyond the scope of this work to dissect the interactions between G_αq/11_, TGFβ2, and PDGF signalling.

As PDGFRβ is a mesenchymal cell marker found outside of the lung, the other organs of *Pdgfrb-Cre^+/−^;Gnaq^fl/fl^;Gna11^−/−^* and *Pdgfrb-Cre/ERT2^+/−^;Gnaq^fl/fl^;Gna11^−/−^* mice were examined histologically. *Pdgfrb-Cre^+/−^;Gnaq^fl/fl^;Gna11^−/−^* kidneys demonstrated expansion and prominence of medullary mesenchymal cells. However, the kidneys of *Pdgfrb-Cre/ERT2^+/−^;Gnaq^fl/fl^;Gna11^−/−^* mice were normal, supporting the hypothesis that abnormalities observed in *Pdgfrb-Cre^+/−^;Gnaq^fl/fl^;Gna11^−/−^* kidneys were developmental in nature.

The limitations of this study predominantly relate to the poor condition of *Pdgfrb-Cre^+/−^;Gnaq^fl/fl^;Gna11^−/−^* mice, which limited the analyses to a single time point and precluded the study of CMS in vivo. Furthermore, the growth restriction of *Pdgfrb-Cre^+/−^;Gnaq^fl/fl^;Gna11^−/−^* mice could have indicated a nutritional deficiency that could have contributed to delayed alveolar development. While these animals did have renal abnormalities which may have contributed to the poor condition and failure to thrive of *Pdgfrb-Cre^+/−^;Gnaq^fl/fl^;Gna11^−/−^* mice, the bowel appeared normal and mice with mesenchymal G_αq/11_ deletion induced in adulthood had normal kidneys. This suggests a true pulmonary phenotype in mesenchymal G_αq/11_ knockout mice. Additionally, our in vitro data provide compelling evidence for a role for mesenchymal G_αq/11_ in a key lung developmental signalling pathway, suggesting that mesenchymal G_αq/11_ deletion generates a true lung developmental phenotype.

Furthermore, while we propose that abnormalities in pericyte differentiation and migration underlie the defective alveologenesis and emphysema in mesenchymal G_αq/11_ knockout mice, *Pdgfrb* is expressed by other cell types, including myofibroblasts, fibroblasts, and vascular smooth muscle cells (Henderson et al. 2013). While it is possible that disturbed TGFβ signalling in these cell types contributed to the lung phenotype in mesenchymal G_αq/11_ knockout mice, pericytes are major progenitors for all these cell types, and are therefore likely to have played a primary role in the abnormalities observed.

Finally, this study has not investigated the role of lung inflammation in mesenchymal G_αq/11_ knockout mice. TGFβ regulates inflammation, and John et al showed that emphysema in mice with a type II epithelial G_αq/11_ deletion was associated with lung inflammation and M2 macrophage polarisation (John *et al*., 2016). The mononuclear cellular aggregates in the lungs of mice with mesenchymal G_αq/11_ deletion induced in adulthood could indicate abnormal inflammation in these mice. However, these cellular aggregates were not observed in mice with a germline mesenchymal G_αq/11_ knockout, and it was not possible to fully define the role of inflammation and the immune response in the emphysema observed in *Pdgfrb-Cre/ERT2^+/−^;Gnaq^fl/fl^;Gna11^−/−^* mice in our study.

In conclusion, this is the first study to generate mesenchymal G_αq/11_ deleted mice, and has demonstrated a novel signalling pathway for CMS-induced TGFβ2 signalling in murine embryonic and mature human mesenchymal cells that is important for alveologenesis and maintenance of the normal lung. These findings could have implications for the treatment of several conditions associated with dysregulated developmental and repair pathways, including fibrosis and emphysema.

## Materials and Methods

### Resource Availability

#### Lead Contact

Further information and requests for resources and reagents should be directed to and will be fulfilled by the Lead Contact, Amanda Goodwin.

#### Materials Availability

This study did not generate new unique reagents.

#### Data and Code Availability

This study did not analyse or generate any new datasets or code

### Experimental Model and Subject Details

#### Animal Studies

##### Husbandry

Mice were housed under specific pathogen-free conditions, with standard food and water available *ad libitum*. All animal experiments were performed in accordance with the Animals (Scientific Procedures) Act 1986, and approved by the Animal Welfare and Ethical Review Board at the University of Nottingham.

##### Breeding strategy

For the germline mouse studies, mice with floxed alleles for *Gnaq* and germline deficiency in *Gna11* (*Gnaq^fl/fl^;Gna11^−/−^*) were crossed with mice that express Cre recombinase under the control of the *Pdgfrb* gene (*Pdgfrb-Cre^+/−^*). *Pdgfrb-Cre^+/−^;Gnaq^+/fl^;Gna11^+/−^* offspring from this F1 generation were then bred with *Gnaq^fl/fl^;Gna11^−/−^* founders to produce an F2 generation, including *Pdgfrb-Cre^+/−^;Gnaq^fl/fl^;Gna11^−/−^* mice. The genetic background for all mice was predominantly C57BL6, with a minimum of a six backcross generations. The generation of *Gnaq^fl/fl^;Gna11^−/−^* and *Pdgfrb-Cre^+/−^* mice has been described previously (Foo et al. 2006; Offermanns et al. 1998; Wettschureck et al. 2001).

For the tamoxifen-inducible mouse gene knockout studies, the same breeding strategy was used as for the germline studies but substituting *Pdgfrb-Cre/ERT2^+/−^* mice (Laboratories) for *Pdgfrb-Cre^+/−^* animals.

##### Genotyping

Mice were genotyped using DNA isolated from ear notch biopsies by PCR analysis with allele-specific primers. Primer sequences: *Cre* transgene 5’- GCG GTC TGG CAG TAA AAA CTA TC – 3’, 5’ - GTG AAA CAG CAT TGC TGT CAC TT – 3’ (product 100bp); internal positive control 5’ - CTA GGC CAC AGA ATT GAA AGA TCT – 3’, 5’ - GTA GGT GGA AAT TCT AGC ATC ATC C – 3’ (product 324bp); *Gna11* wild-type 5’ – AGC ATG CTG TAA GAC CGT AG - 3’, 5’ – GCC CCT TGT ACA GAT GGC AG – 3’ (product 820bp); *Gna11* knockout 5’ - CAG GGG TAG GTG ATG ATT GTG – 3’, 5’ – GAC TAG TGA GAC GTG CTA CTT CC - 3’ (product 450bp); *Gnaq* wild-type and floxed alleles 5’ – GCA TGC GTG TCC TTT ATG TGA G 3’, 5’ – AGC TTA GTC TGG TGA CAG AAG – 3’ (products: 600bp (wild type), 700bp (floxed). For *Cre-ERT2*, the following primers were used: 5’- GAA CTG TCA CCG GGA GGA - 3’, 5’ - AGG CAA ATT TTG GTG TAC GG – 3’ (400bp product).

PCR products were analysed by electrophoresis on ethidium bromide-stained agarose gels.

Mice were genotyped at 2 weeks old (P14). Genotype ratios of F2 mice from the *Gnaq^fl/fl^;Gna11^−/−^* and *Pdgfrb-Cre^+/−^* crosses were compared with the expected Mendelian frequency (12.5% per genotype). Similarly, Genotype ratios of F2 mice from the *Gnaq^fl/fl^;Gna11^−/−^* and *Pdgfrb-Cre/ERT2^+/−^* crosses were assessed, with an expected frequency of 5% for each Cre-expressing genotype.

#### Human Cells

For in vitro experiments using human lung fibroblasts, cells from 4-6 donors were used per group. Cells were used at passage 5-6 for all in vitro experiments.

Human lung fibroblasts (HLFs) were isolated from donated post-mortem or surgical lung biopsy samples, from male and female donors with and without pulmonary fibrosis. For non-fibrotic fibroblasts, cells were isolated from regions of lung distant from the area of primary diagnosis. Tissue was cut into 1mm x 1mm pieces and placed 10mm apart in a 10cm cell culture dish. Tissue was cultured in DMEM supplemented with 10% foetal calf serum (FCS, Fisher), L-glutamine (4mM, Sigma), penicillin (200 units/ml, Sigma), streptomycin (0.2mg/ml, Sigma), and amphotericin B (2.5µg/ml). Fibroblast outgrowth could be seen after 6-8 days. Tissue was removed from the cell culture dish if it became detached, or when cells had reached 80% confluency and were ready for passage. Cells were maintained in a humidified incubator at 37°C, 5% CO_2_/ 95% air, in Dulbecco’s Modified Eagle’s Medium (DMEM, Sigma), supplemented with 10% foetal calf serum (FCS, Fisher), L-glutamine (4mM, Sigma), penicillin (100 units/ml, Sigma) and streptomycin (0.1mg/ml, Sigma).

#### Murine Cells

Wild-type, *Gna12^−/−^;Gna13^−/−^*, and *Gnaq^−/−^;Gna11^−/−^* murine embryonic fibroblasts (MEFs) were a gift from Dr Stefan Offermanns, and their generation has been described elsewhere (Zywietz et al. 2001; Gu et al. 2002). *Gnaq*, *Gna11*, *Gna12*, and *Gna13* gene expression was also confirmed in house prior to these studies. Cells were maintained in a humidified incubator at 37°C, 5% CO_2_/ 95% air, in Dulbecco’s Modified Eagle’s Medium (DMEM, Sigma), supplemented with 10% foetal calf serum (FCS, Fisher), L-glutamine (4mM, Sigma), penicillin (100 units/ml, Sigma) and streptomycin (0.1mg/ml, Sigma).

### Method details

#### Mouse studies

##### Pdgfrb-Cre^+/−^;Gnaq^fl/fl^;Gna11^−/−^ Mouse Phenotyping

Litters were observed for signs of ill health daily from birth. Mice were weighed at P14. Male and female mice were included in all analyses. Mice had not undergone any previous procedures. All mice that survived to P14 were phenotyped and had organs collected. Mouse phenotyping analyses were performed by an observer blinded to genotype. Genotype information was not available to the phenotyping observer until all phenotyping and health status data had been recorded.

##### Tamoxifen-inducible gene knockouts

*Pdgfrb-Cre/ERT2^+/−^;Gnaq^fl/fl^;Gna11^−/−^* offspring and their littermates were kept under standard conditions until 7 weeks of age (P49), when tamoxifen-containing chow (400mg/kg tamoxifen citrate) was introduced *ad libitum*. Health scoring and weights were measured daily for 3 weeks as tamoxifen was administered. Animals were humanely killed after 3 weeks of tamoxifen administration (at 10 weeks old, P70).

##### Organ Collection

Mice were humanely killed by intraperitoneal injection of pentobarbital, and organs collected for histological analyses. The lungs were perfused by injecting 40units/ml heparin sodium in PBS (Wockhardt) into the right ventricle, and inflated by cannulating the trachea and filling the lungs with 10% formalin (VWR) under gravity. The trachea was ligated, and the heart and lungs removed en bloc. Livers and kidneys were also collected. Organs were kept in 10% formalin (VWR) for 24 hours before paraffin embedding and sectioning.

#### Tissue histology staining

3µm (lung, kidney), and 5µm (heart, liver) formalin-fixed paraffin embedded tissue sections were deparaffinised in xylene and rehydrated in graded alcohols. Haematoxylin and eosin, Verhoeff van Gieson (elastin), and picrosirius red staining were performed as per standard protocols using buffers and stains prepared in house and mounted in DPX.

#### Staining solutions made in house

The following histology solutions were generated in house: Weigert’s iodine (2g potassium iodide, 1g iodine, 100ml distilled water); Verheoff’s solution (20ml 5% alcoholic haematoxylin, 8ml 10% ferric chloride, 8ml Weigert’s iodine); Van Gieson’s solution (5ml aqueous acid fuschin, 100ml saturated aqueous picric acid); Picro-sirius red solution (0.5g Direct Red 80 (Sigma), 500ml saturated aqueous picric acid); Weigert’s haematoxylin (1:1 ratio of Weigert’s solution A and Weigert’s solution B); Weigert’s solution A (1% haematoxylin in 100% ethanol); Weigert’s solution B (4ml 30% ferric chloride, 1ml 12N hydrochloric acid, 95ml water); Acidified water (5ml glacial acetic acid, 1l distilled water); Acid/alcohol solution (70% ethanol, 0.1% hydrochloric acid).

##### Haematoxylin and eosin (H&E) stain

After being deparaffinised and rehydrated, tissue sections were submerged in Mayers haematoxylin (Fisher) for 2 minutes, acid/alcohol solution for 1 minute, then 1% eosin solution (VWR) for 3 minutes. Sections were rinsed with tap water between each step, then dehydrated and mounted.

##### Elastin (Verhoeff Van Gieson) stain

Lung sections were deparaffinised and hydrated to distilled water, then stained in Verhoeff’s solution for 1 hour until the tissue was completely black. Sections were differentiated in 2% ferric chloride until elastin fibres were seen on a grey background, incubated in 5% sodium thiosulphate for 1 minute, and then washed in running tap water for 5 minutes. Sections were then counterstained in Van Gieson’s solution for 5 minutes, dehydrated and mounted as above.

##### Picrosirius red stain

Lung, kidney, and heart sections were deparaffinised and hydrated. Nuclei were stained with Weigert’s haematoxylin for 8 minutes, and then washed in running tap water for 5 minutes. Sections were incubated in picrosirius red for 1 hour, washed in two changed of acidified water, then dehydrated and mounted.

#### Immunostaining

Tissue sections were deparaffinised in xylene and rehydrated in graded alcohols. Heat-mediated antigen retrieval was performed by boiling sections in a microwave for 20 minutes in 10mM citric acid buffer (pH 6.0). Endogenous peroxidase activity was blocked by incubating sections in 3% hydrogen peroxide in methanol for 30 minutes. Nonspecific binding was blocked with 5% goat serum (Sigma) in 0.1% BSA/PBS. Sections were incubated with primary antibody in 5% goat serum overnight at 4°C in a humidified chamber, followed by incubations for 60 minutes with secondary antibody and 30 minutes with avidin-biotin complex (Vector). Sections were then stained with diaminobenzidine (Sigma), counterstained with Mayers haematoxylin (Sigma), and mounted in DPX (Sigma). Slides were washed in PBS (Sigma) between incubation steps.

The following antibodies were used for immunohistochemistry: Rabbit anti-αSMA (Abcam, ab5694; 1:500), rabbit anti-CD31 (Abcam, ab182981; 1:2000), rabbit anti-ki67 (Abcam, ab15580; 1µg/ml), rabbit anti-pro-surfactant protein C (Sigma, Ab3786; 1:2000), rabbit anti-TGFβ2 (Proteintech, 19999-1-AP; 1:3000), rabbit anti-elastin (Atlas, HPA056941; 1:100), and biotinylated goat anti-rabbit IgG (Vector, BA1000; 1:200).

#### Image Quantification

##### Image acquisition

Images of H&E, elastin, and IHC were taken using a Nikon 90i microscope and NIS-Elements software v3.2 (Nikon). Polarised light imaging of picrosirius red stained samples was performed using a Zeiss Axioplan microscope (Zeiss) and MicroManager 1.4 software (Vale Lab, UCSF).

##### Staining quantification

For all analyses of histology images, *Pdgfrb-Cre^+/−^;Gnaq^fl/fl^;Gna11^−/−^* or *Pdgfrb-Cre/ERT2^+/−^;Gnaq^fl/fl^;Gna11^−/−^* mice were compared with *Pdgfrb-Cre^−/−^;Gnaq^fl/fl^;Gna11^−/−^* littermate controls (labelled as *Gna11^−/−^* controls). For histological analyses, four animals per genotype were assessed to allow differences in histological appearances to be detected. All image quantification was performed by an observer blinded to genotype. This observer was not unblinded to genotype until all image quantification data had been recorded.

For quantitative analyses of the lungs, 5-10 images were assessed per set of lungs, covering all lobes and avoiding major airways and central blood vessels. All morphometric analyses were performed using NIS Elements software v3.2 (Nikon), with the exception of peripheral pulmonary vessel thickness measurements and kidney measurements, which were performed using CaseViewer 2.3 software (3D Histech).

For quantification of immunohistochemistry and elastin staining, the “count” feature of ImageJ (NIH) was used. Elastin fibres were identified as thin black fibres on VVG stain, and secondary crests were elastin positive if they had black staining that was not clearly a cell nucleus on Verhoeff van Geison staining. For immunohistochemistry staining, a cell was counted if it stained brown. Only nuclear DAB staining was counted for Ki67 quantification. For αSMA quantification, the number of αSMA-positive secondary crests per 40 x field was counted. For Ki67 and pro-SPC staining, the total number of cells per 40x field was quantified by counting nuclei, and the proportion of Ki67 or pro-SPC positive cells calculated by dividing the number of stained cells per image by the total number of cells per image.

For quantification of TGFβ2 staining, the following scoring system was used and 7 fields (20x magnification) per mouse were analysed:

- **Score 0**: No cells stained.
- **Score 0.5**: 1-25 cells stained at low intensity
- **Score 1.0**: 1-25 cells stained at high intensity
- **Score 1.5**: 26-50 cells stained at low intensity
- **Score 2.0**: 26-50 cells stained at high intensity
- **Score 2.5**: >50 cells stained at low intensity
- **Score 3.0**: >50 cells stained at high intensity

##### Morphometry

Mean linear intercept (MLI) analysis of airspace size was performed as previously described (John et al. 2016). Briefly, 10x magnification images were overlaid with a grid comprised of 100µm squares, and “intercepts” between gridlines and airspace walls counted. The MLI was calculated by dividing the length of each gridline was divided by the intercept count. For alveolar wall thickness measurements, 40x magnification images were overlaid with five equally spaced horizontal lines and the alveolar wall thickness measured at points where lung tissue crossed each line using the “measure” function of NIS Elements. Mean MLI and alveolar wall thickness values were calculated for each mouse from all measurements across all images and data presented as median ± interquartile range. For secondary crest counts, 10x magnification images were used and secondary crests counted for each image. For peripheral vessel wall thickness, ten random peripheral pulmonary vessels were identified using CD31 staining. Maximal and minimum vessel wall thickness in µm was measured using the “measure” function of CaseViewer. For assessment of right ventricular hypertrophy, the left and right cardiac ventricular wall thickness was measured using CaseViewer, and the right: left ventricular wall thickness ratio calculated.

#### Breathing-related cyclical stretch experiments

Cells were seeded at 2 x 10^5^ cells per well on collagen I-coated Bioflex® 6 well culture plates (Dunn Labortechnik) in DMEM supplemented with 10% FCS, L-glutamine (4mM), penicillin (100 units/ml) and streptomycin (0.1mg/ml) and allowed to adhere for 24 hours. The culture medium was changed to 1% FCS in DMEM with 4mM L-glutamine for 24 hours before stretching commenced. The Flexcell® FX-5000T system (Flexcell International Corporation) was used to apply cyclical stretch to cells in vitro, according to the manufacturer’s instructions. MEFs were stretched at a frequency of 1Hz, and HLFs at 0.3Hz to mimic breathing in the relevant organism. 15% elongation and a sine waveform were used for all cyclical stretch experiments. Cyclical stretch was applied for 48 hours, except for experiments using siRNA-induced *GNAQ* and *GNA11* knockdown, where 24 hours of cyclical stretch was used. Unstretched control cells were cultured in identical conditions alongside the Flexcell® apparatus. Cells were lysed in protein lysis buffer (Cell Signalling) supplemented with phosphatase (Phos-Stop, Sigma) and protease (Complete Mini, Sigma) inhibitors, and 20µM PMSF. All experimental replicates were performed independently.

##### Chemical Inhibitors used in Cyclical Stretch System

When used, inhibitor compounds were applied in DMEM supplemented with 1% FCS and 4mM L-glutamine 30 minutes before stretching commenced. The activin receptor-like kinase (ALK5)/ type I TGFβ-receptor kinase inhibitor SB-525334 (Sigma) was used at a concentration of 50µM. A ROCK inhibitor (Y27632, Sigma), pan-αv integrin inhibitor (CWHM-12), β1 integrin inhibitor (NOTT199SS) matrix metalloproteinase (MMP) inhibitor GM6001 (Sigma), and serine protease inhibitor AEBSF (Sigma) were used at varying concentrations. Where inhibitors were dissolved in DMSO, the negative control cells were treated with a DMSO concentration equivalent to that used in the highest inhibitor concentration.

##### GNAQ and GNA11 siRNA

SiRNAs for human *GNAQ* (Dharmacon ON-TARGET-plus SMARTpool GNAQ) and *GNA11* (Dharmacon ON-TARGET-plus SMARTpool GNA11) were used to induce *GNAQ* and *GNA11* knockdown. A non-targeting siRNA pool was used as a control (Dharmacon ON-TARGET-plus non-targeting pool).

Cells were seeded at 1.5 x 10^5^ cells per well of a 6 well Flexcell® plate in antibiotic-free DMEM supplemented with 10% FCS and 4mM L-glutamine. The following day, *GNAQ* and *GNA11* siRNA was applied at a concentration of 15nM each with 4µl/ml DharmaFECT 1 transfection reagent (Dharmacon) as per the manufacturer’s protocol. At 48 hours after transfection, the media was changed to DMEM supplemented with 1% FCS and 4mM L-glutamine. Cyclical stretch was applied for 24 hours from 72 hours post-transfection. LPA stimulation was applied for 4 hours from 72 hours post-transfection. G_αq/11_ knockdown was confirmed by western blot and qPCR

#### Western blotting

Protein concentrations were determined by BCA assay using a commercially available kit (ThermoFisher), according to the manufacturer’s instructions. Equal amounts of protein (15-25µg) were loaded per lane of a 10% SDS-polyacrylamide gel and subject to electrophoresis, and transferred onto a polyvinylidene fluoride membrane (BioRad). Membranes were blocked for 1 hour in either 5% non-fat milk (pSmad2, Smad2/3, αSMA, G_αq_, G_α11_, GAPDH) or 3% BSA (TGFβ1, TGFβ2) in tris-buffered saline containing 0.1% Tween, pH 7.4 (TBST). Membranes were incubated overnight at 4°C in blocking buffer with the appropriate primary antibody. Membranes were washed in TBST, then incubated for 1-2 hours in the appropriate HRP-conjugated secondary antibody in blocking buffer. Western blots were analysed using chemilluminescence and exposure to film (GE Healthcare). Where membranes were probed for two different proteins of the same molecular weight, i.e. pSmad2 and Smad2, the membrane was stripped after analysis of pSmad2 using Western Restore Stripping Buffer (Thermo-Fisher) for 5 minutes and re-blocked with 5% non-fat milk before application of the second primary antibody.

The following antibodies were used for western blots: Rabbit anti-phospho-Smad2 (pSmad2) (Cell Signaling Technology, 3808; 1:1000), rabbit anti-Smad2/3 (Cell Signaling Technology, 3102; 1:1000), rabbit anti-αSMA (Abcam, ab5694; 0.5µg/ml), rabbit anti-GAPDH (Abcam, ab181603; 1:10,000), rabbit anti-TGFβ1 (ab92486; 4µg/ml), mouse anti-TGFβ2 (Abcam, ab36495; 1:1000), rabbit anti G_α11_ (Abcam, ab153951; 1:1000), goat anti-G_αq_ (Abcam, ab128060; 0.1µg/ml), HRP-conjugated goat-anti-rabbit (Agilent, P044801-2; 1:3000), HRP-conjugated rabbit-anti-goat (Agilent, P016002-2; 1:3000), HRP-conjugated rabbit anti-mouse (Agilent, P0260022-2, 1:3000).

##### Densitometry Analysis of Western Blots

Densitometry was performed using ImageJ (NIH) on scanned western blot images. JPEG images were converted into greyscale images, and the software used to calculate densitometry values for each band relative to the other bands. These relative densitometry values were used to calculate the expression of protein relative to loading control using the equation: Protein relative to loading control = protein densitometry value/ loading control protein densitometry value

#### Quantitative PCR

RNA was isolated from in vitro experiments using the Machery-Nagel Nucleospin RNA isolation kit according to the manufacturer’s instructions. Complementary DNA (cDNA) was reverse transcribed from 200µg RNA using Superscript IV Reverse Transcriptase (Thermo Fisher) according to the manufacturer’s protocol. Quantitative PCR was performed on cDNA using gene-specific primers (see below), and an MXPro3000 qPCR machine (Stratagene) at an annealing temperature of 60°C for 40 cycles. KAPA SYBR FastTaq (Sigma) was used for qPCR of all genes other than *Pdgfa*, *Pdgfb*, *Pdgfc*, *Pdgfd*, *Pdgfra*, and *Pdgfrb*, for which PerfeCTa SYBR Green Fastmix (VWR) was used. Amplification of a single PCR product was confirmed by melting curve analysis. The delta-delta Ct method was used to quantify gene expression relative to the housekeeping genes *Hprt* (mouse samples) or *B2M* (human samples).

Primer sequences for mouse genes were: *Hprt* forward 5’ – TGA AAG ACT TGC TCG AGA TGT CA - 3’, *Hprt* reverse 5’ CCA GCA GGT CAG CAA AGA ACT 3’, *Acta2* forward 5’ GGG ATC CTG ACG CTG AAG TA 3’, *Acta2* reverse 5’ GAC AGC ACA GCC TGA ATA GC 3’, *Eln* forward 5’ GAT GGT GCA CAC CTT TGT TG 3’, *Eln* reverse 5’ CAG TGT GAG CCA TCT CA 3’, *Col1a1* forward 5’ AGC TTT GTG CAC CTC CGG CT 3’, *Col1a1* reverse 5’ ACA CAG CCG TGC CAT TGT GG 3’, *Col3a1* forward 5’ TTT GCA GCC TGG GCT CAT TT 3’,*Col3a1* reverse 5’ AGG TAC CGA TTT GAA CAG ACT, *Pdgfa* forward 5’ GAG ATA CCC CGG GAG TTG A 3’, *Pdgfa* reverse 5’ TCT TGC AAA CTG CAG GAA TG 3’, *Pdgfb* forward 5’ TGA AAT GCT GAG CGA CCA C 3’, *Pdgfb* reverse 5’ AGC TTT CCA ACT CGA CTC C 3’, *Pdgfc* forward 5’ AGG TTG TCT CCT GGT CAA GC 3’, *Pdgfc* reverse 5’ CCT GCG TTT CCT CTA CAC AC 3’, *Pdgfd* forward 5’CCA AGG AAC CTG CTT CTG AC 3’, *Pdgfd* reverse 5’ CTT GGA GGG ATC TCC TTG TG 3’, *Pdgfra* forward 5’ CAA ACC CTG AGA CCA CAA TG 3’, *Pdgfra* reverse 5’ TCC CCC AAC AGT AAC CCA AG 3’, *Pdgfrb* forward TGC CTC AGC CAA ATG TCA CC 3’, *Pdgfrb* reverse 5’ TGC TCA CCA CCT CGT ATT CC 3’.

Primer sequences for human genes were: *GNAQ* forward 5’ – GGACAGGAGAGGGTGGCAAG – 3’, *GNAQ* reverse 5’ – TGGGATCTTGAGTGTGTCCA – 3’, *GNA11* forward 5’ – CCACTGCTTTGAGAACGTGA – 3’, *GNA11* reverse 5’ GCAGGTCCTTCTTGTTGAGG – 3’, *B2M* forward 5’AATCCAAATGCGGCATCT3’, *B2M* reverse 5’GAGTATGCCTGCCGTGTG3’.

#### Statistical Analyses

Statistical analyses were performed using GraphPad Prism 8.2 software (GraphPad). For experiments with group sizes of 5 or less, a non-parametric test was used. For experiments with group sizes of 6 or over, data were assessed for normality and a parametric test used if data followed a normal distribution.

#### Key Resources

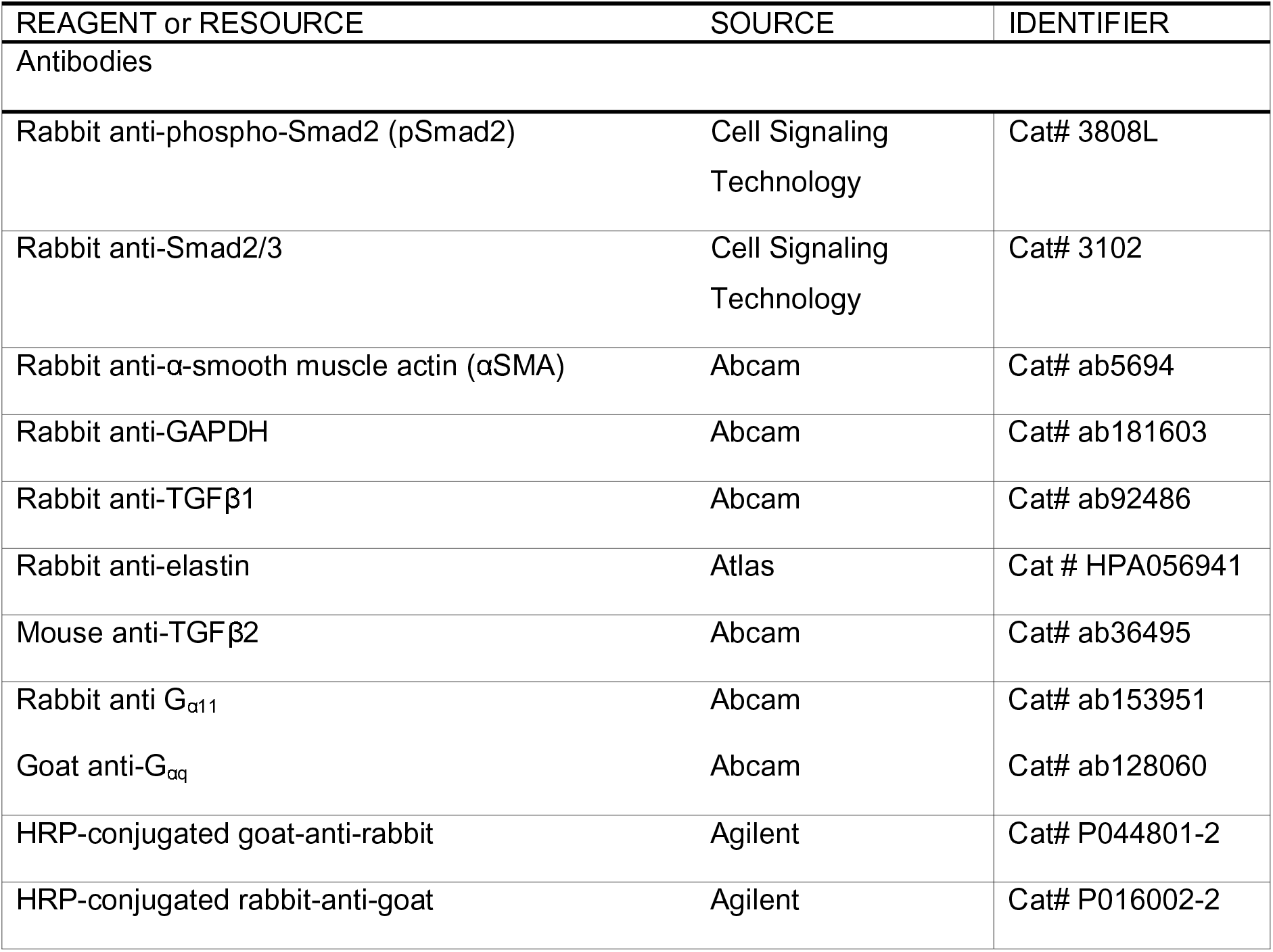
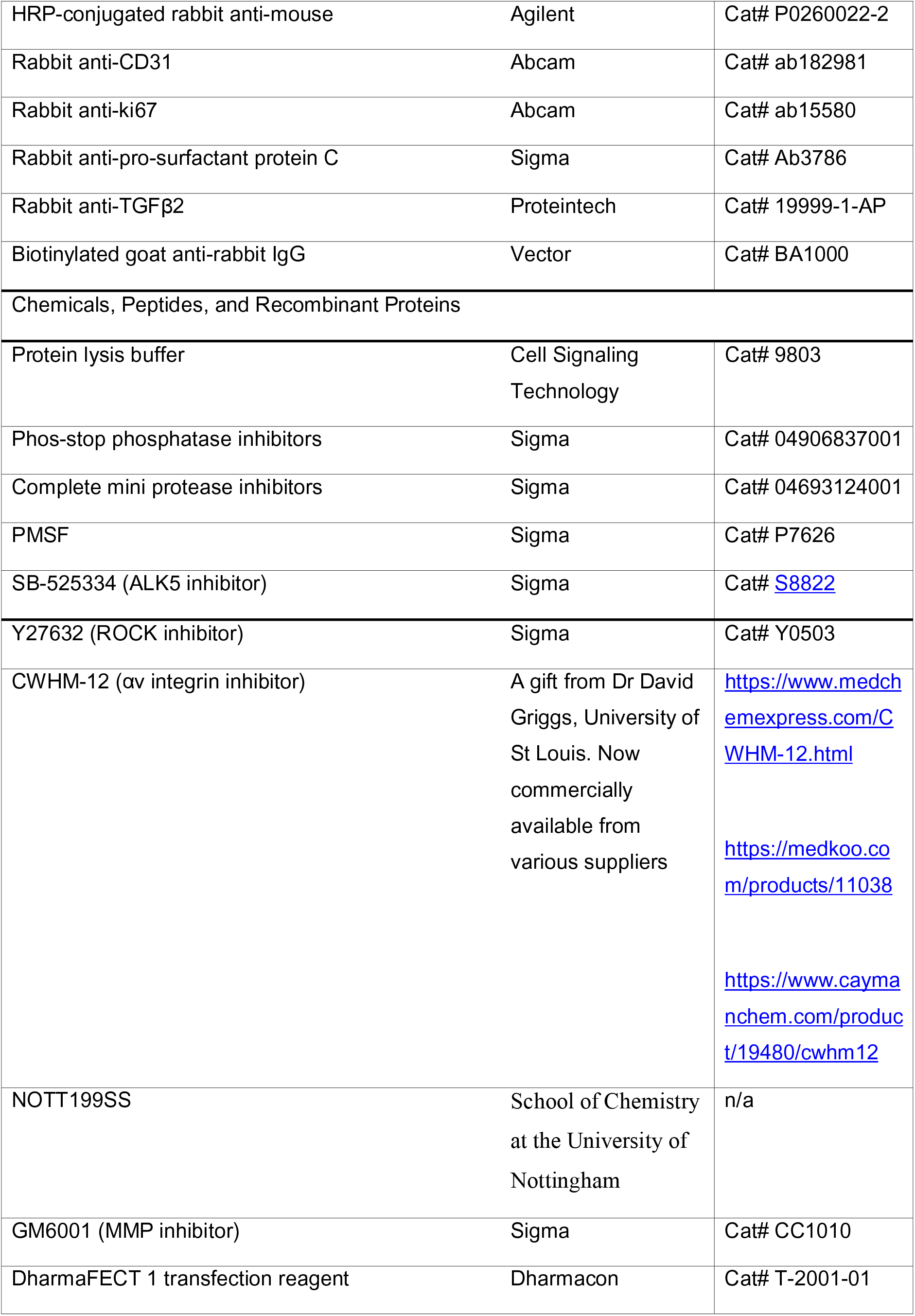
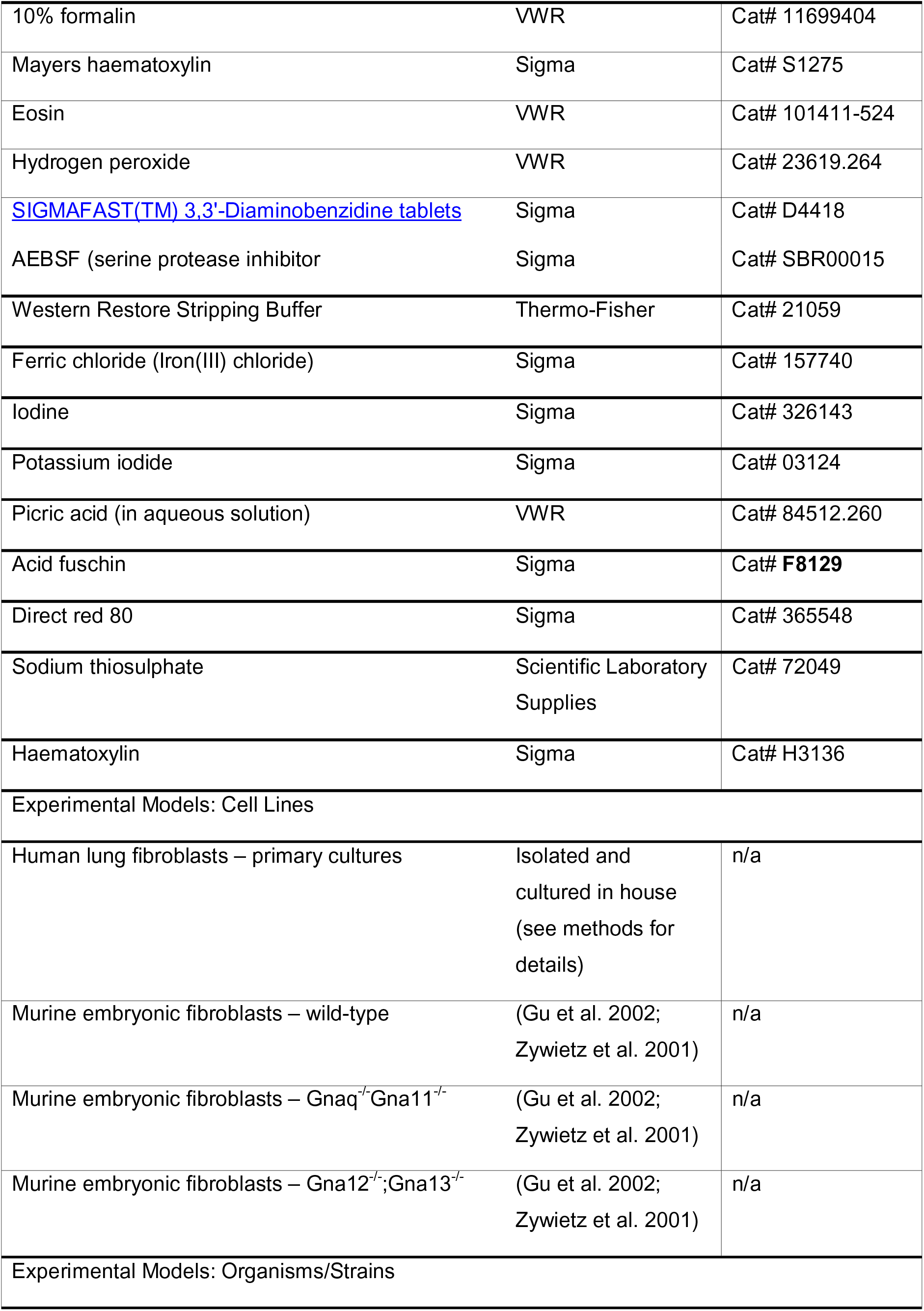
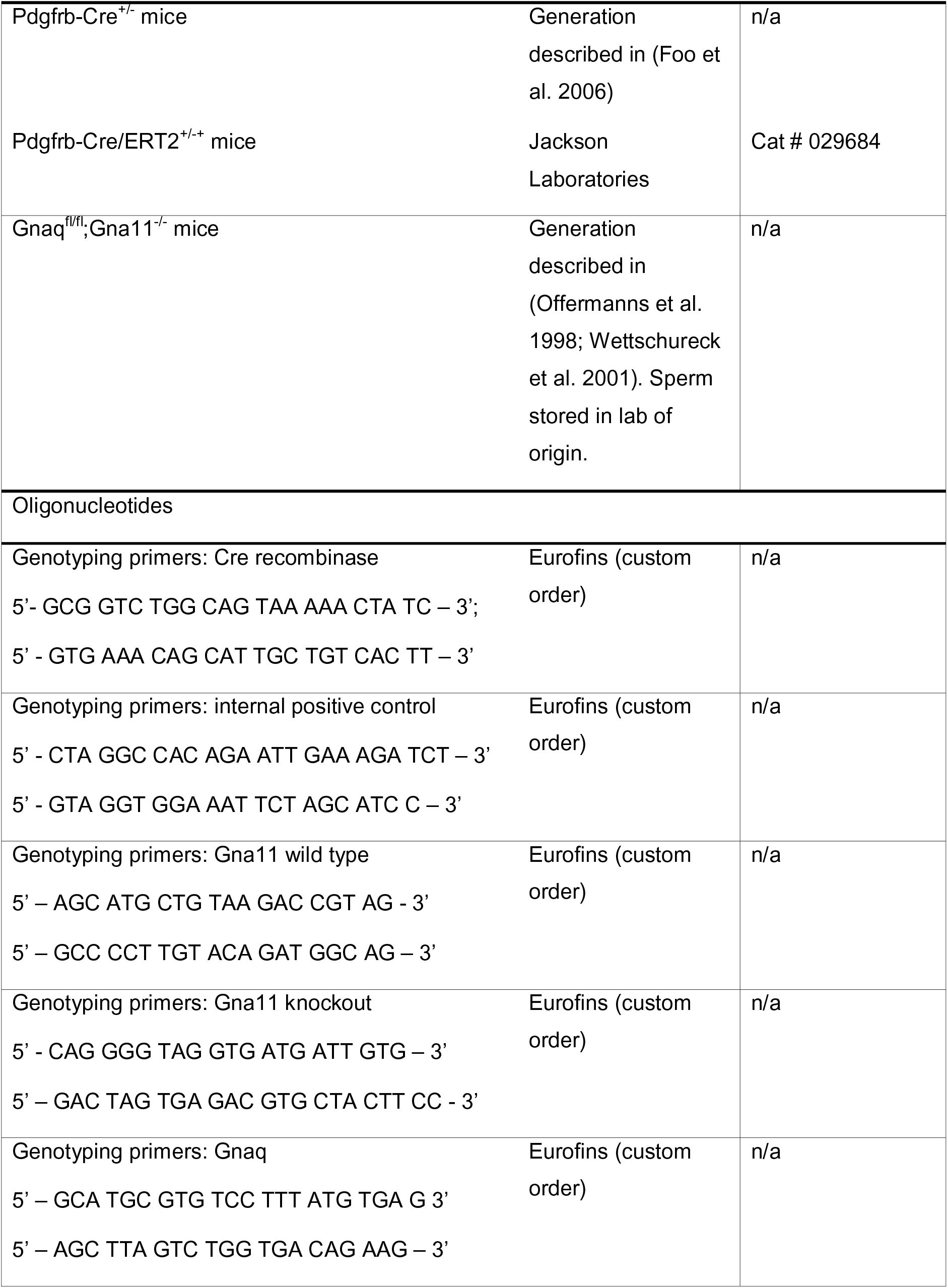
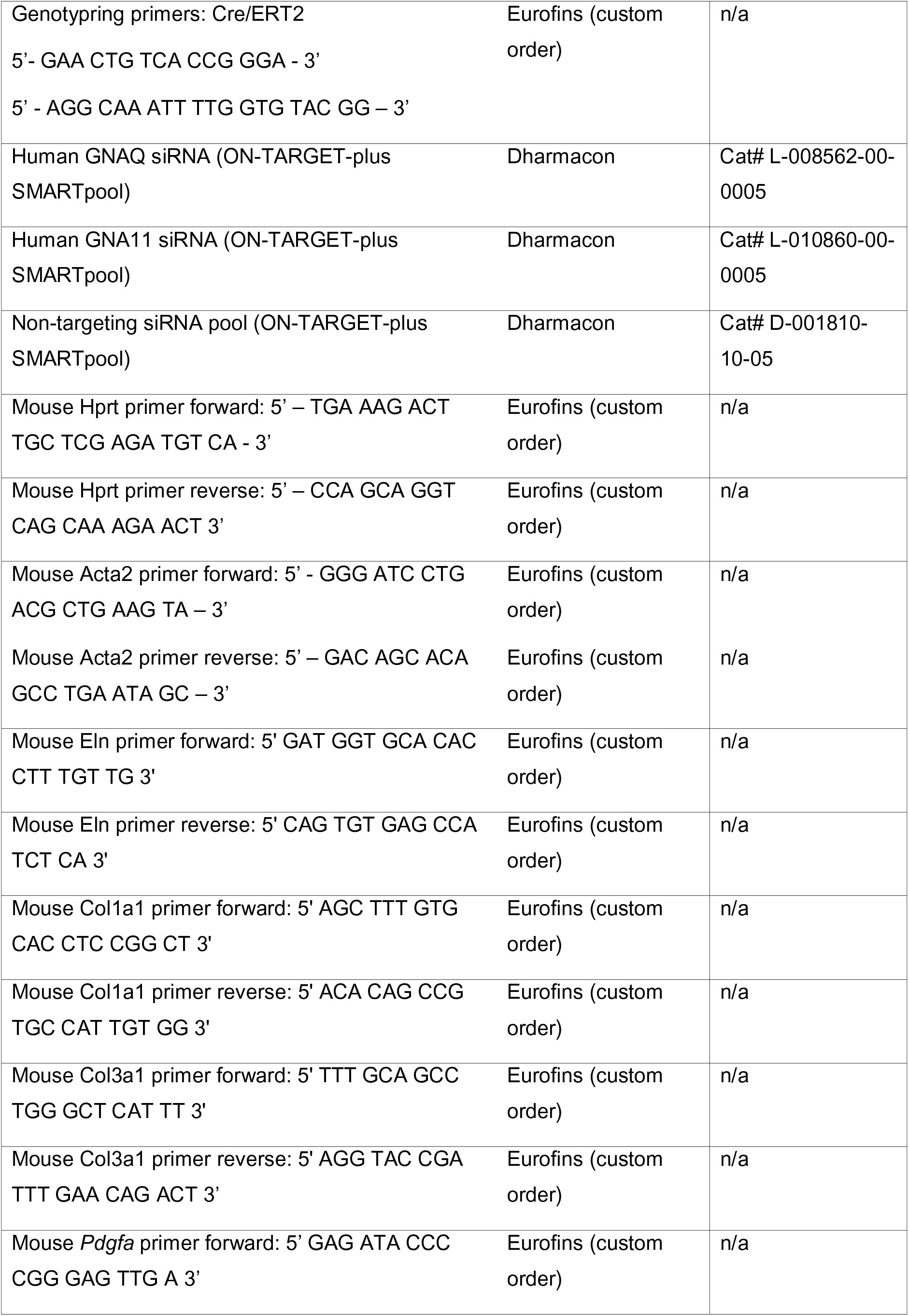
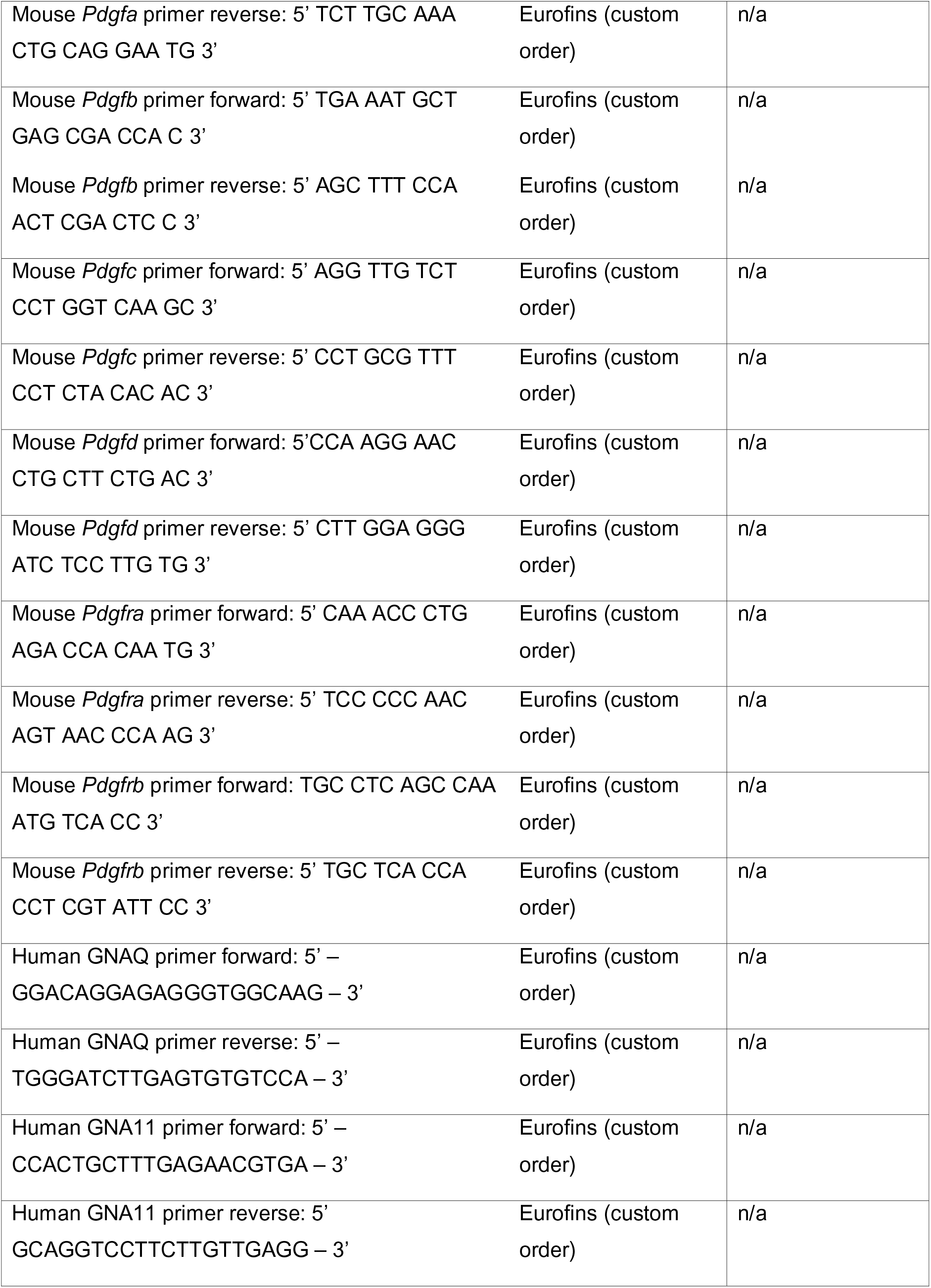
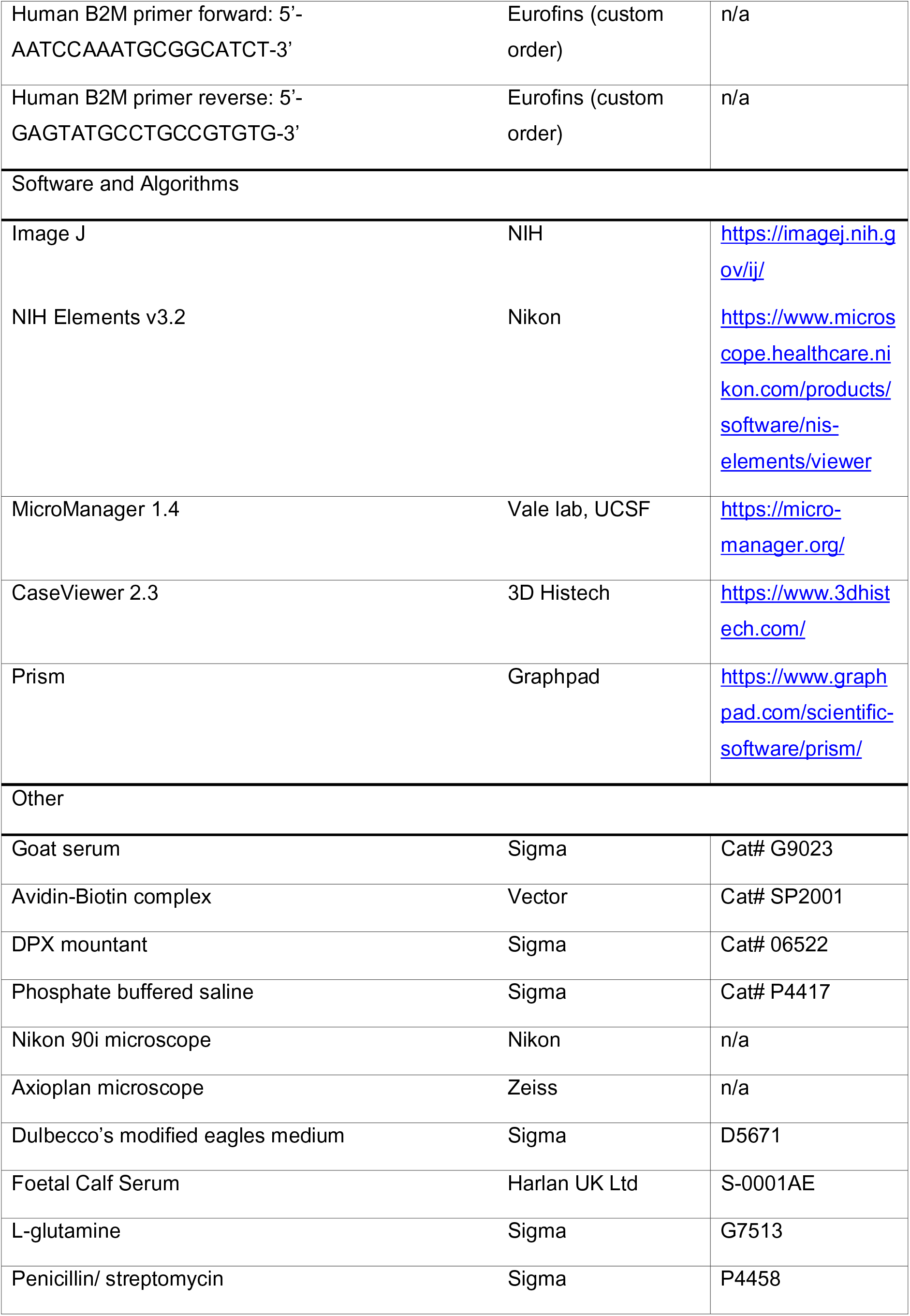
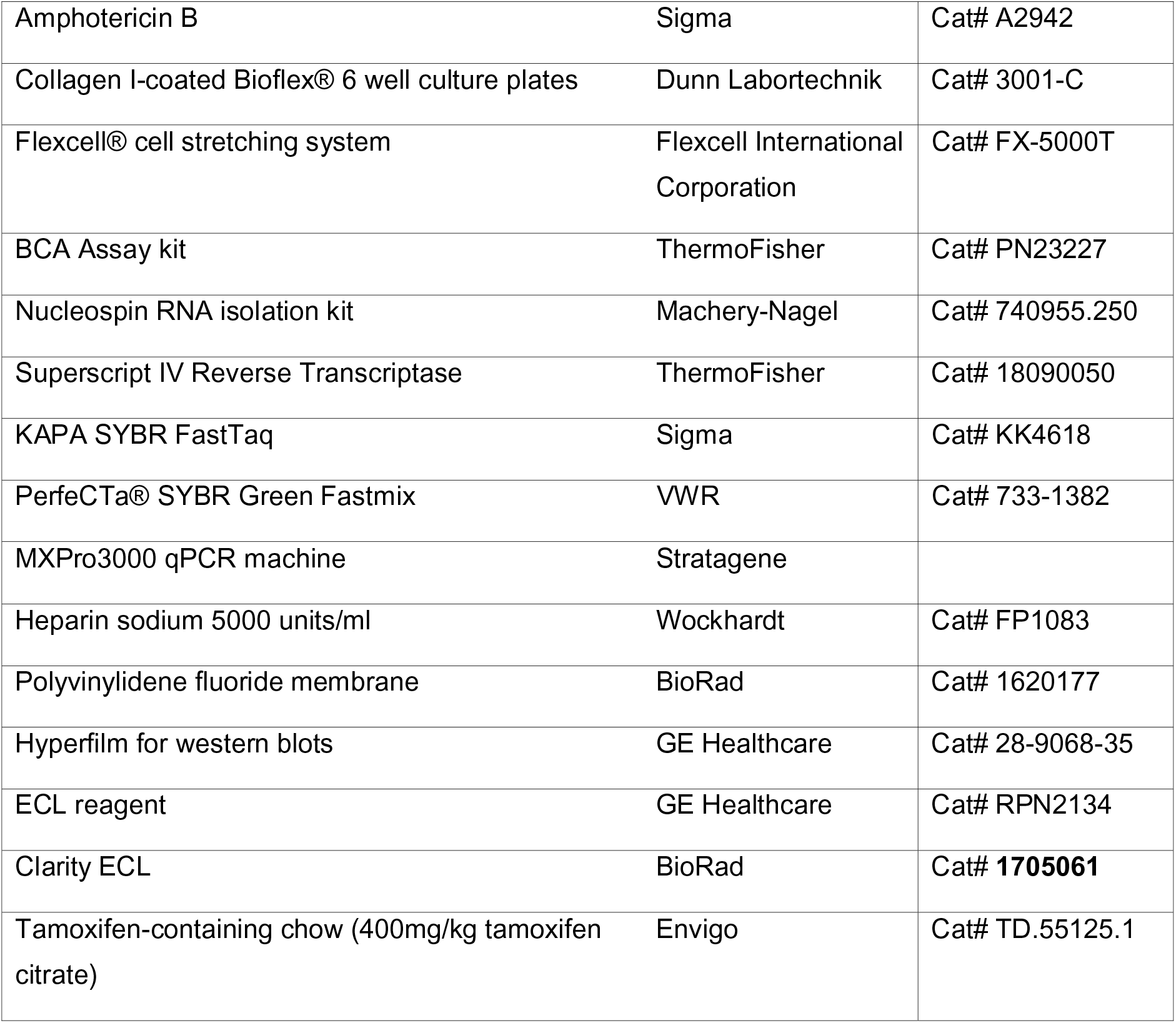

## Acknowledgements

We thank Dr Thomas McInally (University of Nottingham, UK) and Dr David Griggs (St Louis University, Missouri, USA) for supplying the compounds NOTT199SS and CWHM-12, respectively. NOTT199SS was identified as part of an MSci Chemistry undergraduate integrin drug discovery collaboration between the School of Chemistry at the University of Nottingham and GlaxoSmithKline (GSK), supervised by Dr Simon Macdonald (GSK) and Dr Thomas McInally (University of Nottingham). We also thank Dr Tim Kendall (University of Edinburgh), for his opinion on the liver histology.

## Author contributions

ATG and GJ conceived project. ATG performed experiments, conducted image analysis, and wrote the original draft manuscript. AEJ supervised and gave methodological guidance for mouse phenotyping experiments and microscopy. CJ performed some of the immunohistochemistry included in this work. ATG, AEJ, AH, and ALT conducted animal monitoring and tissue collection. ALT established original cultures of human lung fibroblasts used in this study. KS and MP provided expert guidance on the performing and interpretation of kidney histology. NCH and SO provided and guided the breeding of *Pdgfrb-Cre^+/−^* and *Gnaq^fl/fl^;Gna11^−/−^* mice, respectively. GJ supervised the entire project. All authors reviewed the original draft manuscript and contributed to editing and preparation of the final manuscript.

## Supplemental Information titles and legends

Figure S1: *Pdgfrb-Cre^+/−^;Gnaq^fl/fl^;Gna11^−/−^* mice have normal liver, heart, and bowel histology.

Representative images of haematoxylin and eosin staining of liver (A), heart (B), and bowel (C) from *Gna11^−/−^* and *Pdgfrb-Cre^+/−^;Gnaq^fl/fl^;Gna11^−/−^* mice.

Figure S2: *Pdgfrb-Cre/ERT2^+/−^;Gnaq^fl/fl^;Gna11^−/−^* mice have normal kidney histology after three weeks of tamoxifen.

Representative images of haematoxylin and eosin staining of kidney (A) from *Gna11^−/−^* and *Pdgfrb-Cre/ERT2^+/−^;Gnaq^fl/fl^;Gna11^−/−^* mice treated with three weeks of tamoxifen.

Figure S3: Cyclical stretch-induced TGFβ activation occurs independently of ROCK, and αv and β1 integrins in fibroblasts.

A) Representative pSmad2 western blot of human lung fibroblasts treated with Y27632 then subject to 48 hours of cyclical stretch (15% elongation, 0.3Hz, 48 hours).

B) Relative pSmad2 to Smad2 densitometry from western blots of human lung fibroblasts treated with Y27632 then subject to cyclical stretch. Median ± interquartile range, n=4, two-tailed Mann Whitney test.

C) Representative pSmad2 western blot of human lung fibroblasts treated with an αv integrin inhibitor (CWHM-12) then subject to 48 hours of cyclical stretch (15% elongation, 0.3Hz, 48 hours).

D) Relative pSmad2 to Smad2 densitometry of human lung fibroblasts treated with CWHM-12 then subject to cyclical stretch. Median ± interquartile range, n=4, two-tailed Mann Whitney test.

E) Representative pSmad2 western blot of human lung fibroblasts treated with a β1 integrin inhibitor (NOTT199SS) then subject to 48 hours of cyclical stretch (15% elongation, 0.3Hz, 48 hours).

F) pSmad2 relative to Smad2 densitometry of human lung fibroblasts treated with NOTT199SS then subject to cyclical stretch. Median ± interquartile range, n=4, two-tailed Mann Whitney Test.

+ = stretched; - = unstretched. Alk5 inh = 50µM Alk5 inhibitor (SB525334)

