## Supplementary figures and images for "Stretch Regulates Alveologenesis and Homeostasis Via Mesenchymal G_αq/11_-Mediated TGFβ2 Activation"

### Supp figure 1

**Supp Fig 1**

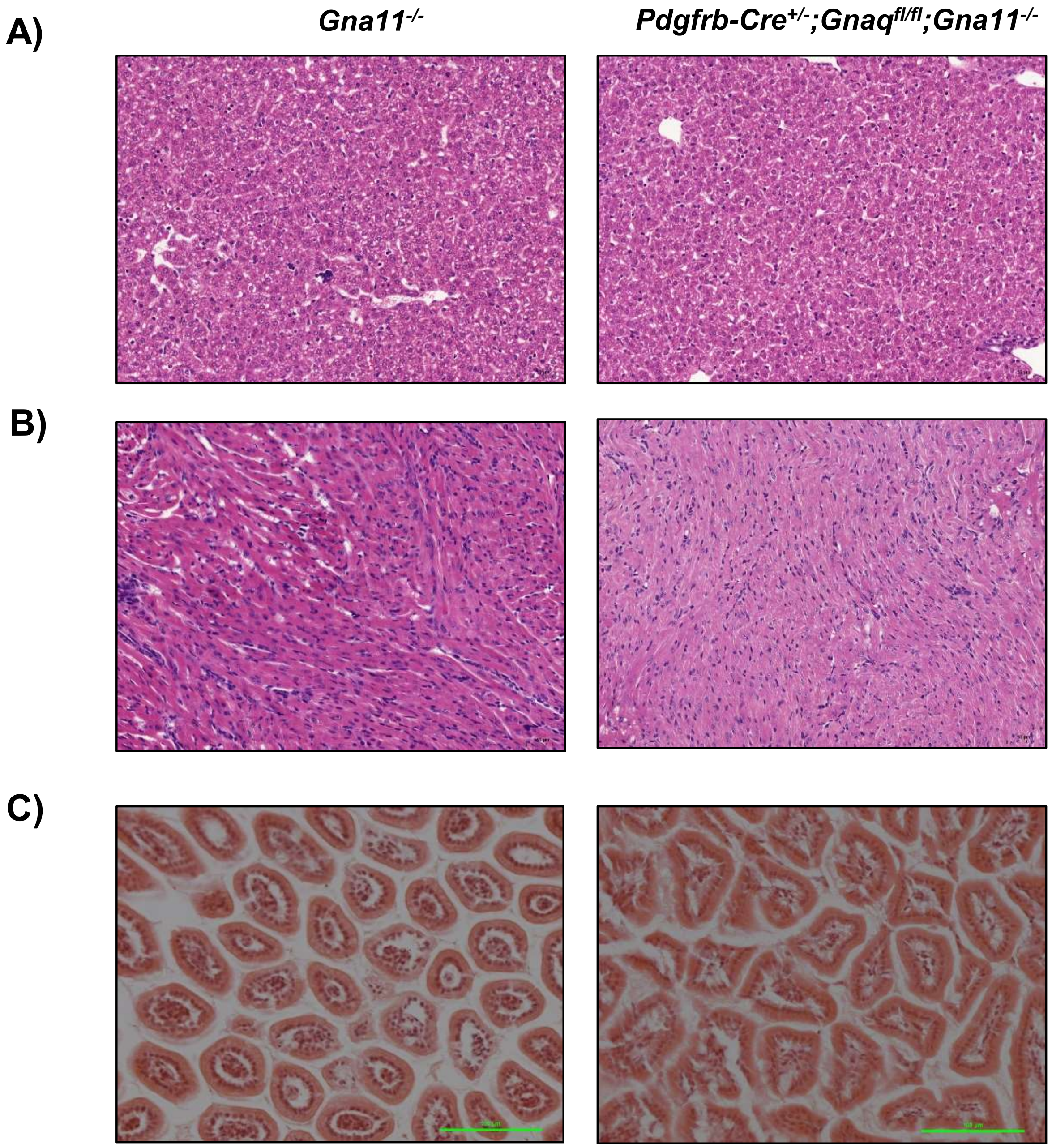

### Supp figure 2

#### Supp Fig 2

**A)**

*Gna11* $^{-/-}$ *Pdgfrb-Cre/ERT2* $^{+/-}$ *;Gnaq* $^{fl/fl}$ *;Gna11* $^{-/-}$

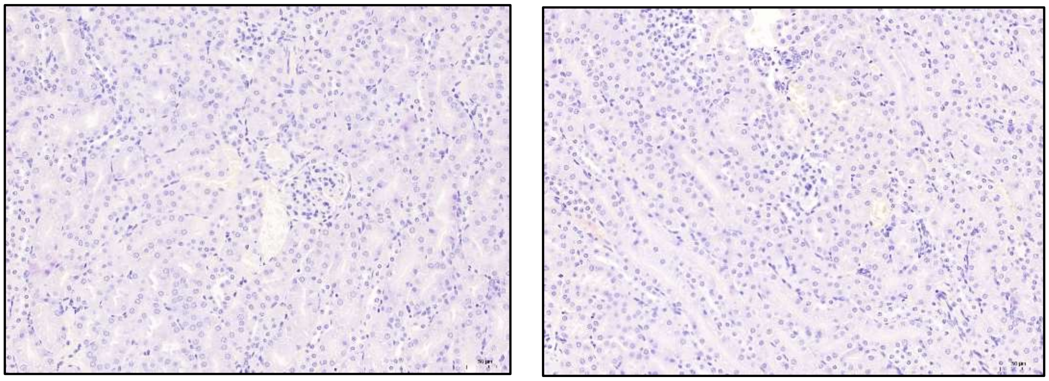

### Supp figure 3

#### Supp figure 3

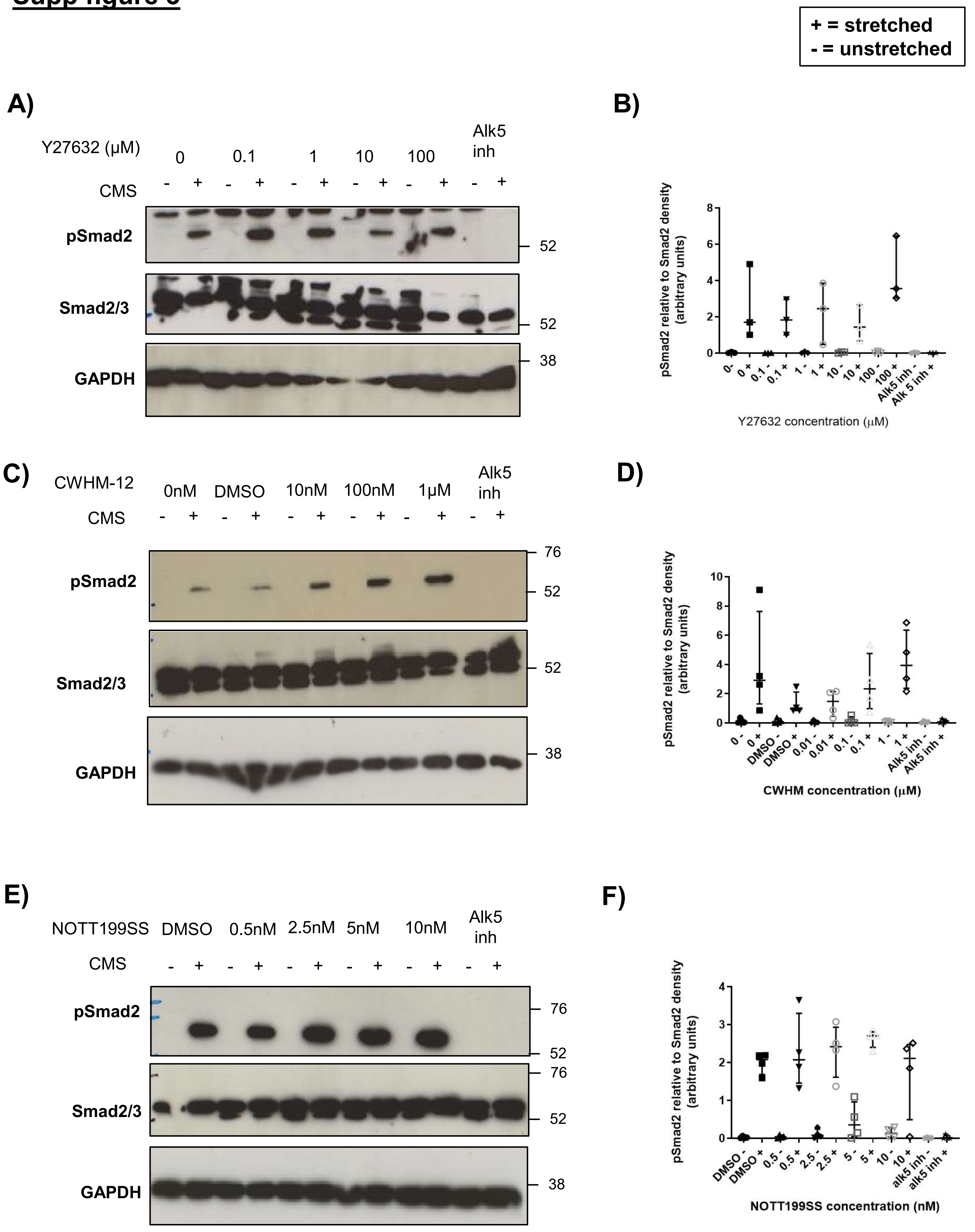
